# Striatal pathways oppositely shift cortical activity along the decision axis

**DOI:** 10.1101/2025.07.29.667406

**Authors:** Jounhong Ryan Cho, Scott S. Bolkan, Lindsey S. Brown, Maja Skuza, Yousuf El-Jayyousi, Benjamin Midler, Robert N. Fetcho, Christopher A. Zimmerman, Alejandro Pan-Vazquez, Manuel Schottdorf, Adrian G. Bondy, Misael A. Sanchez, Juan F. Lopez Luna, Alvaro Luna, Tim Eilers, Abigail S. Kalmbach, Yang Lu, Laura A. Lynch, Ilana B. Witten

## Abstract

The cortex and basal ganglia are organized into multiple parallel loops that serve motor, limbic, and cognitive functions. The classic model of cortico-basal ganglia interactions posits that within each loop, the direct pathway of the basal ganglia activates the cortex and the indirect pathway inhibits it^1–3^. While this model has found support in the motor domain^4,5^, whether opponent control by the two pathways extends to the cognitive domain remains unknown. Here, we record from anterior cingulate cortex (ACC) and dorsomedial striatum (DMS) while inhibiting direct or indirect pathway neurons in DMS, as mice perform an accumulation-of-evidence task^6–10^. Inconsistent with the classic model, the manipulations do not produce opponent changes in overall ACC activity. Instead, the pathways exert opponent influence over a subpopulation of ACC neurons that encode accumulated sensory evidence, the task-relevant decision variable. The direction of the modulation depends on a neuron’s tuning to ipsilateral versus contralateral evidence, such that the two pathways generate opponent shifts in coding specifically along the decision axis. Thus, our results uncover unexpected specificity in the effects of basal ganglia pathways on the cortex, with the two pathways of the DMS exerting opponent control not on overall activity but on coding of the relevant task variable. This functional specificity may extend to other basal ganglia loops to support different aspects of adaptive behavior, with the pathways serving a general role in selecting and shifting cortical representations to subserve circuit-specific functions.

## Main

The direct pathway of the basal ganglia has been associated with promoting movement and activating the motor cortex. In contrast, the indirect pathway, via divergent projection patterns (**Fig. 1a**), is thought to have an opponent role in suppressing movement and motor cortical activity^1–3^. This framework has proven valuable for understanding movement disorders^11–16^, and is supported by animal studies that focus on motor control^4,5,17–26^. However, if and how this framework extends to non-motor functions of the basal ganglia remains unknown.

**Fig. 1:**
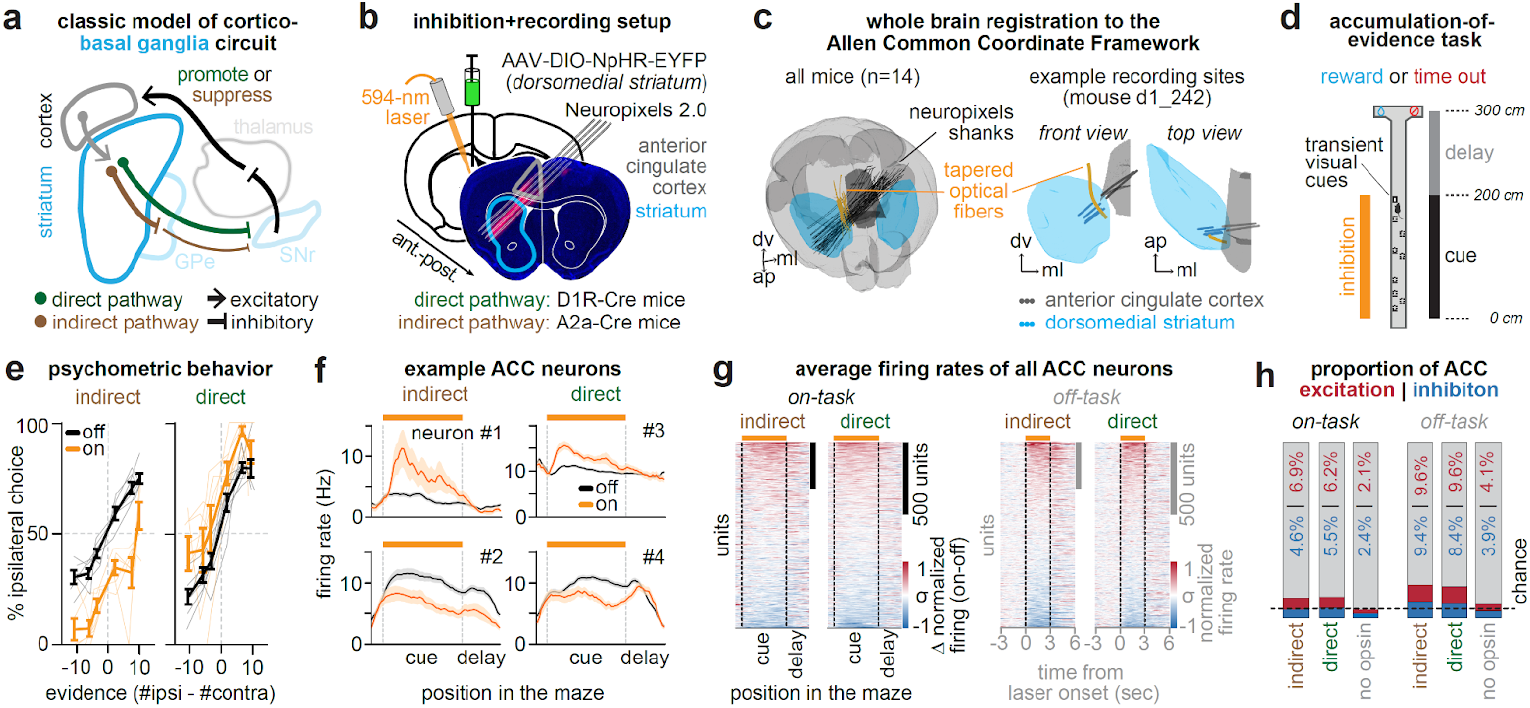
Striatal pathway inhibition has opposing influence over decisions but not overall ACC activity. **a**, Schematic of the cortico-basal ganglia-thalamic loop. **b**, Schematic of experimental setup: Cre-dependent viral expression of halorhodopsin (NpHR) and a tapered optical fiber were targeted to the dorsomedial striatum (DMS) of the right hemisphere in D1R-Cre (direct pathway) or A2a-Cre (indirect pathway) mice; Neuropixels 2.0 probes were targeted to the anterior cingulate cortex (ACC) and striatum of the same hemisphere. **c**, *Left*, Registration to the Allen Common Coordinate Framework (CCF) of all Neuropixels 2.0 shank and optical fiber trajectories (n = 14 mice). *Right*, Registration of fiberoptic and recording sites targeting ACC (grey) or DMS (blue) for an example mouse. **d**, Schematic of the virtual reality based accumulation-of-evidence task. Laser was delivered to the DMS during the cue region on a random subset (10-20%) of trials. **e**, Behavioral performance during the task as a function of the difference in visual cues relative to the laser hemisphere (#ipsilateral - #contralateral) during laser off (black) or on (orange) trials, shown separately for mice receiving indirect (*left*, n = 6 mice) or direct (*right*, n = 6 mice) pathway inhibition. Thick lines: mean across mice. Transparent lines: mean of individual mice. **f**, Position-binned activity (Hz) during the task for four example ACC neurons averaged across laser off (black) or on (orange) trials. Example neurons are from mice receiving indirect (*left column*) or direct (*right column*) pathway inhibition. **g**, Trial averaged activity for all ACC neurons during the task (*left two plots*) or off-task (*right two plots*), with indirect (n = 1698 neurons from 12 sessions) or direct (n = 1220 neurons from 15 sessions) pathway inhibition. Activity is z-score normalized and sorted by the magnitude of laser modulation in each task condition and inhibition group. **h**, Proportion of significantly excited (red) or inhibited (blue) ACC neurons as a function of task (*left three bars*: on-task; *right three bars*: off-task) and inhibition group (indirect pathway; direct pathway; or, no opsin control: n = 573 neurons from 2 mice across 5 sessions). Data are presented as mean ± s.e.m.

In particular, basal ganglia loops interconnected to the frontal cortex are thought to contribute to cognition rather than motor control^27–34^. For example, transient inhibition of the two pathways within the dorsomedial striatum (DMS), which forms a loop with the anterior cingulate cortex (ACC)^35,36,23,37^, has little impact on spontaneous movements or sensory-guided movements^9,38^. In contrast, the same manipulation produces large and opponent behavioral effects when mice make decisions based on the gradual accumulation of sensory evidence^9^.

How do DMS pathways generate opponent control that is specific to the decision-making process? Here, we investigate the hypothesis that opponent control of behavior by DMS pathways is mediated in part through selective effects on decision-related representations in the cortex. To test this idea, we characterize how cortical activity is affected by inhibition of each pathway during an evidence accumulation task in which the two pathways have opposing effects on behavior.

### Combining large-scale ACC and DMS recordings with DMS pathway inhibition

We performed chronic Neuropixels 2.0 recordings in mice across ACC and DMS of the same hemisphere, while unilaterally inhibiting the striatal indirect or direct pathway (**Fig. 1b-c**; indirect: n = 2,030 ACC & 1,618 DMS single- and multi-unit neurons from 6 mice; direct: n = 1,449 ACC & 1,319 DMS neurons from 6 mice). We focused on the DMS and ACC because they form part of an anatomically verified cortico-basal ganglia loop^23,37^. To inhibit each pathway, we used Cre-dependent viral expression of halorhodopsin^39^ in the DMS of A2a-Cre mice to target the indirect pathway, and D1R-Cre mice to target the direct pathway (**Fig. 1b**), as previously validated^9^.

Recording site locations and fiber tracks were verified using whole-brain clearing, light-sheet microscopy, and registration of fiber and electrode tracks to the Allen Common Coordinate Framework based on electrophysiological landmarks^40,41^ (**Fig. 1c**; **Extended Data Fig. 1a-b**). To reduce the influence of duplicate neurons when sampling the same recording site across multiple days, we used the software package UnitMatch^42^ to remove clusters based on metrics of waveform similarity (see **Methods, Neuropixels Data Acquisition**, *Removal of duplicate clusters*).

### Opposing influence of DMS pathways on decisions but not overall ACC activity

To assess the influence of DMS pathways on behavior and ACC activity, we optogenetically inhibited each pathway and recorded neural activity while head-fixed mice ran on a spherical treadmill in the dark (“off task”, **Extended Data Fig. 2a**), or performed an accumulation-of-evidence task^6,43,7,8,10^ (“on task”, **Fig. 1d**).

In the accumulation-of-evidence task, mice navigated a T-maze in virtual reality while transient visual cues were presented along both maze walls. After a delay, mice were rewarded for turning to the side that had the greater number of cues throughout the maze. Thus, the task requires the accumulation of sensory evidence as new visual inputs arrive, and the transformation of this memory into a choice. Pathway-specific inhibition was delivered unilaterally on a random subset of trials (10-20%) and was limited to the cue region of the task (**Fig. 1d**).

Consistent with the two DMS pathways exerting opponent control over evidence-guided decision-making^9^, unilateral indirect pathway inhibition during the task produced a decision bias ipsilateral to the laser hemisphere (**Fig. 1e**, *left*) while direct pathway inhibition produced a contralateral bias (**Fig. 1e**, *right*). This effect was strongest when mice were task-engaged, as assessed with a GLM-HMM behavioral model^9,44^ (**Extended Data Fig. 3**).

According to the classic cortico-basal ganglia model (**Fig. 1a**), indirect and direct pathway inhibition should increase and decrease cortical activity, respectively. Inconsistent with this prediction, ACC neurons displayed heterogeneous responses to pathway-specific inhibition, with both excitation and inhibition observed following either manipulation (example neurons, **Fig. 1f**; all neurons, **Fig. 1g**). This was true whether pathway inhibition occurred “on task” or “off task”.

Overall, we observed a tendency towards greater ACC excitation over inhibition when inhibiting either pathway (**Fig. 1h**, *left*; excitation vs inhibition “on task”: indirect, 117 vs 78/1698 neurons, p<0.01; direct, 73 vs 64/1174 neurons, p=0.43; **Fig. 1h**, *right*, “off task”: indirect, 163 vs 159/1698 neurons, p=0.82; direct, 113 vs 99/1174 neurons, p=0.31; two-tailed 2-proportion Z-test). These effects were not explained by the influence of tissue heating on neural activity^45,46^, as control mice without opsin expression had significantly fewer light-modulated neurons compared to mice expressing halorhodopsin (**Fig. 1h**; **Extended Data Fig. 2c**). Consistent with prior work^4^, there were also fewer light-modulated neurons in the “on task” compared to “off task” setting (**Fig. 1h**, “on task” vs “off task”: indirect: 195 vs 322/1698 neurons; direct: 137 vs 212/1174 neurons; p<0.0001, two-tailed 2-proportion Z-test), potentially because task demands recruit cortical activity that is less susceptible to pathway perturbations.

Together, these results suggest that the classic model in which the two basal ganglia pathways exert opponent control over behavior via broadly promoting and suppressing cortical activity is incomplete.

### Identification of task-coding neurons

Given that inhibition of DMS pathways oppositely influences decision-making (**Fig. 1e**), but did not have opposite effects on overall ACC activity (**Fig. 1g-h**), we wondered if DMS pathways may selectively influence the subset of ACC neurons encoding the task. To test this, we isolated two populations of task-related ACC neurons: one which encoded graded levels of sensory evidence (“evidenced-tuned”), and another with binary encoding of the selected choice (“choice-tuned”)^47^. In this section, we describe the identification and characterization of these decision-related neurons in ACC and DMS, and in the next sections the effect of pathway inhibition on their firing patterns.

To identify these neurons, on laser off trials, we fit a linear encoding model (see **Methods, Neural Analyses**, *Linear encoding model*) for each neuron at each maze position to predict neural activity across trials based on the accumulated sensory evidence, the behavioral choice, as well as the previous trial outcome (**Extended Data Fig. 4**). Neurons were defined as ‘evidenced-tuned’ if they showed significance in the evidence coefficient at any position in the cue and delay regions (**Fig. 1d**), and ‘choice-tuned’ if they showed significance for the choice coefficient but not the evidence coefficient (see **Methods, Neural Analyses**, *Statistical definition of task-relevant neurons*).

Evidence-tuned neurons showed graded changes in firing rates as a function of accumulated evidence (example neurons: **Fig. 2a**, *top row*, and **Extended Data Fig. 4d,g**; population averaged ACC activity: **Fig. 2b**, *top row*; population averaged striatal activity: **Extended Data Fig. 5**), with preferences (i.e. increased firing rate) for either ipsilateral or contralateral evidence (relative to the recording hemisphere). This dependence of activity on evidence was largely preserved on incorrect trials (example neuron: **Fig. 2a**, *top right*; population averaged ACC activity: **Fig. 2b**, *top row*). Evidence-tuned activity rose during the cue region when visual cues were presented and persisted into the delay region prior to choice execution (**Fig. 2c**, *top row*). ACC and dorsal striatum contained similar proportions of neurons significantly tuned to evidence (**Fig. 2d**; ACC vs striatum: 462/3147 vs 274/2136 neurons, p=0.06).

**Fig. 2:**
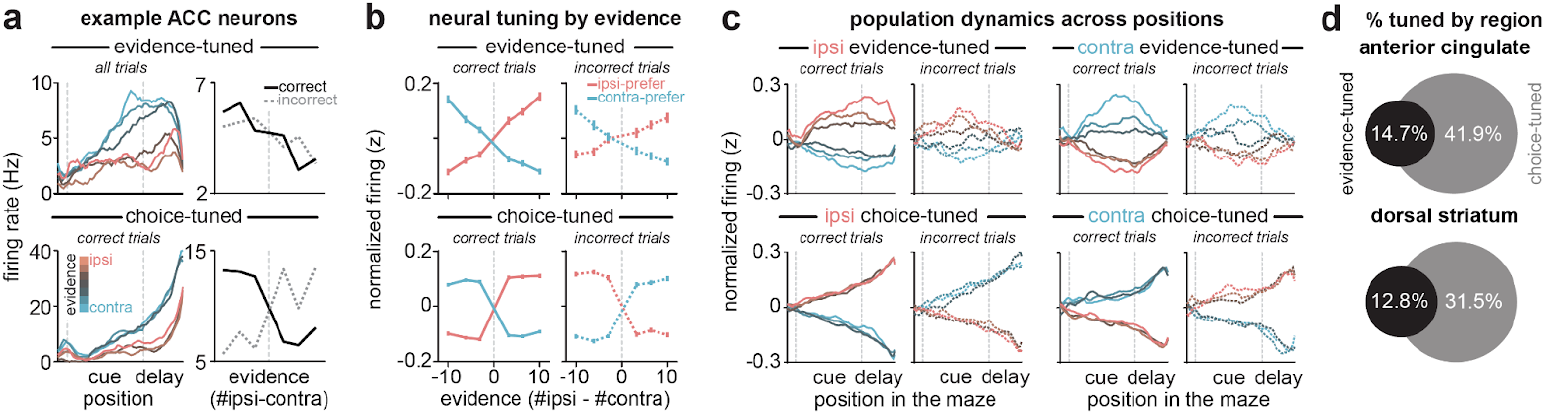
ACC neurons encode decisions with graded tuning to sensory evidence and/or binary tuning to the behavioral choice. **a**, Example ACC neurons with significant tuning to evidence (*top row*) or choice (*bottom row*). *Left column*: Position-binned firing rate (Hz) in the task, averaged across trials binned by sensory evidence level (#ipsilateral-#contralateral cues). *Right column*: Evidence tuning curves, constructed by averaging neural activity (Hz) in the cue and delay regions across trials binned by sensory evidence level (at the end of the maze). Averages were taken across correct (black) and incorrect (dotted gray) trials separately. Only correct trials were included for the example choice-tuned neuron (*bottom left*) due to the inverted firing rates observed on incorrect trials (*bottom right*). **b**, Evidence tuning curves, as in **a**, *right column*, but for z-scored activity averaged across all evidence-(*top row*; n = 224 ipsilateral- and 238 contralateral-preferring), or choice-tuned ACC neurons (*bottom row*; n = 693 ipsilateral- and 624 contralateral-preferring), parsed by their ipsilateral (salmon) or contralateral (aqua) preference. Averaged activity on correct (*left column*; solid lines) and incorrect (*right column*; dotted lines) trials shown separately. **c**, Averaged z-scored activity across position in the task for all evidence-(*top row*) or choice-tuned (*bottom row*) ACC neurons parsed by their ipsilateral (*left two columns*) or contralateral (*right two columns*) preference, and displayed separately for correct (solid line) and incorrect (dashed line) trials. **d**, Proportion of ACC (*top*) and dorsal striatal (*bottom*) neurons with significant tuning to either evidence (black) or choice but not evidence (grey) (ACC: n = 3147 total neurons from 27 sessions; dorsal striatum: n = 2136 total neurons from 23 sessions). Data are presented as mean in **a** and **c**, and mean ± s.e.m. in **b**.

In contrast to evidence-tuned neurons, choice-tuned neurons encoded the sign of accumulated evidence on correct trials, and reversed their evidence preference on incorrect trials (example neurons: **Fig. 2a**, *bottom row*, and **Extended Data Fig. 4e,h**; population averaged ACC activity: **Fig. 2b**, *bottom row*; population averaged striatal activity: **Extended Data Fig. 5**). This is consistent with coding of the behavioral choice (ipsilateral or contralateral) rather than the evidence. Choice-tuned activity was present throughout the task, but was most prominent during the delay region prior to choice execution (example neuron: **Fig. 2a**, *bottom right*; population averaged ACC activity: **Fig. 2c**, *bottom row*). While dorsal striatum contained many choice-tuned neurons, there was a greater proportion in ACC (**Fig. 2d;** ACC vs striatum: 1317/3147 vs 672/2136 neurons; p<0.0001, two-tailed 2-proportion Z-test).

### Indirect pathway inhibition shifts ACC activity towards contralateral evidence representations

We next sought to determine the effects of pathway inhibition on these subpopulations of decision-related neurons in ACC. To this end, we re-fit the linear encoding model including an additional regressor for laser delivery (**Fig. 3a;** see **Methods, Neural Analyses**, *Linear encoding model*), and examined laser coefficients and evidence tuning curves in neurons classified by their functional tuning profile. We first examined how indirect pathway inhibition affected evidence-tuned neurons.

**Fig. 3:**
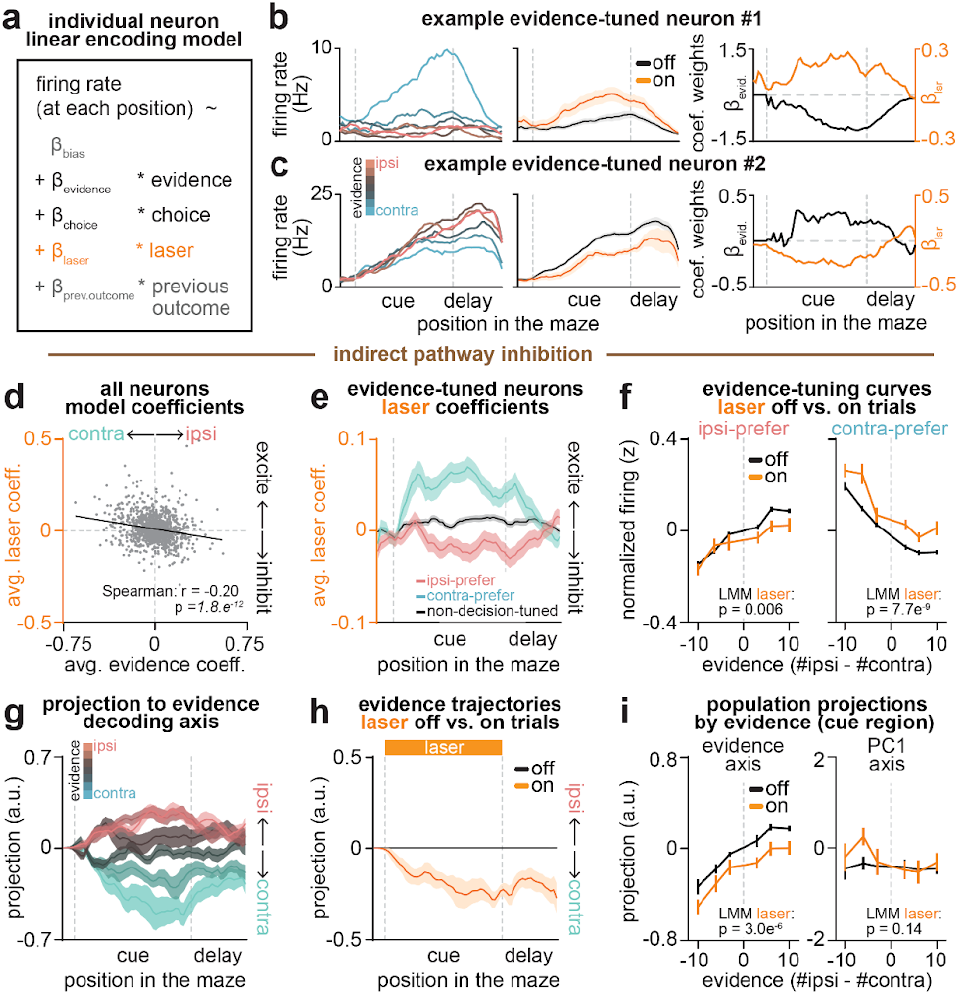
Indirect pathway inhibition shifts ACC activity towards contralateral-evidence representations. **a**, Linear encoding model used to predict the activity of each neuron at each position based on task variables. All correct/incorrect and laser on/off trials were used to fit the model. **b**, An example ACC neuron with indirect pathway inhibition that showed significance for evidence (negative coefficient: contralateral preference) and laser (positive coefficient: excitation). *Left*: Mean position-binned firing rate (Hz) averaged across trials binned by sensory evidence level (#ipsilateral - #contralateral cues). *Middle*: Same as *left* but averaged across all laser off (black) or on (orange) trials. *Right*: Model coefficients for evidence (black) and laser (orange) across positions. **c**, Same as **b**, but for an ACC neuron with significant tuning to evidence (with positive coefficient: ipsilateral preference) and laser (with negative coefficient: inhibition). **d**, Average evidence vs. laser coefficients across cue and delay positions in the task for all ACC neurons in indirect pathway mice (grey dots; n = 2030 neurons from 12 sessions). Black line: Spearman rank correlation; r = −0.20; p < 0.0001. **e**, Averaged laser coefficients at each position in the task for all ACC neurons that showed significant tuning for ipsilateral evidence (salmon; n = 136 neurons), contralateral evidence (aqua; n = 140), or neurons without significant tuning to evidence or choice (black, n = 938). **f**, Evidence tuning curves for all ACC neurons with significant evidence-tuning, parsed by ipsilateral-(*left*) or contralateral (*right*) preference. For each neuron, z-scored activity was averaged in all cue and delay positions across trials binned by sensory evidence level, separately for laser off (black) or on (orange) trials. Statistics reflect significance of a linear mixed-effects model (ipsilateral: p_laser_<0.01; contralateral: p_laser_<0.0001). **g**, Cross-validated projection of ACC population activity onto a one-dimensional evidence decoding axis at each position in the maze (n = 12 sessions with more than 50 neurons), and then averaged across laser off trials binned by sensory evidence level. **h**, Same as **g**, but for the averaged evidence-axis projection across held-out laser off (black) or on (orange) trials. Positive (or negative) values (a.u., arbitrary unit) indicate greater ipsilateral (or contralateral) evidence representation. **i**, Similar to **f**, but evidence tuning curves from population activity projected onto the evidence-axis (*left*) or first principle component (PC1)-axis (*right*). Tuning curves were constructed by averaging projected population activity across positions in the cue region and trials binned by sensory evidence level (#ipsilateral - #contralateral cues), separately for laser off (black) or on (orange) trials. Statistics reflect significance of a linear mixed-effects model (evidence axis: p_laser_<0.0001; PC1 axis: p_laser_=0.14). Data are presented as mean ± s.e.m. unless otherwise stated.

Even when considering only the subpopulation of evidence-tuned neurons, indirect pathway inhibition produced heterogeneous effects on ACC activity. For instance, the contralateral-preferring evidence neuron shown in **Fig. 3b** was excited by indirect pathway inhibition, while the ipsilateral-preferring one in **Fig. 3c** was inhibited by the same manipulation.

However, despite this heterogeneity, there was a compelling pattern across the subpopulation of evidence-tuned ACC neurons. Consistent with the two example neurons (**Fig. 3b-c**), indirect pathway inhibition tended to excite (positive laser coefficient) neurons that preferred contralateral evidence, and inhibit (negative laser coefficient) neurons that preferred ipsilateral evidence (**Fig. 3d-e**).

To visualize this effect directly in the ACC firing rates, we plotted population-averaged evidence tuning curves separately for ipsilateral- and contralateral-preferring evidence-tuned neurons, both with and without indirect pathway inhibition (**Fig. 3f**). Consistent with the encoding model coefficients (**Fig. 3d-e**), indirect pathway inhibition decreased the activity of ipsilateral-preferring neurons, and increased that of contralateral-preferring neurons (**Fig. 3f**).

In theory, the net effect of inhibiting ipsilateral- and exciting contralateral-preferring evidence-tuned neurons would be to shift the population activity towards representing contralateral evidence. To test this, we identified the dimension of ACC population activity along which evidence was best decoded at each maze position (“decision axis”; **Extended Data Fig. 6a**; see **Methods, Neural Analyses**, *Obtaining evidence decoder axis*). As expected based on how this dimension was defined, projecting neural activity from laser off trials onto this axis at each position revealed clear separation of neural trajectories across trials with varying levels of evidence (**Fig. 3g**). Similar to the evidence tuning curves (**Fig. 2b**, *top*), the evidence separation was similarly present on both correct and incorrect trials (**Extended Data Fig. 6c**). Comparing the projection from held-out trials with versus without laser delivery revealed a shift towards greater contralateral evidence coding with indirect pathway inhibition (**Fig. 3h**). This shift was apparent across evidence levels (**Fig. 3i**, *left*).

This net shift towards contralateral evidence coding in ACC with indirect pathway inhibition is consistent with the contralateral decision bias in the behavior (**Fig. 1e**), suggesting a potential causal link. Indeed, at the level of individual sessions, the extent of laser-induced shifts in evidence representations correlated with the degree of behavioral bias (**Extended Data Fig. 6g**).

In contrast to the decision axis, neural projections onto the first principal component axis – which primarily captures the spatio-temporal progression of mice across the maze (**Extended Data Fig. 6d-f**) – showed little modulation by indirect pathway inhibition (**Fig. 3i**, *right*; **Extended Data Fig. 6e**). This suggests specificity of the effect of indirect pathway inhibition on the decision axis in ACC.

### Direct pathway inhibition shifts ACC activity towards ipsilateral evidence representations

We next asked how evidence-tuned ACC neurons were impacted by direct pathway inhibition. In contrast to the indirect pathway (**Fig. 3**), direct pathway inhibition increased activity of ipsilateral-, but not contralateral-preferring, evidence-tuned neurons (encoding model coefficients: **Fig. 4a,b**; population-averaged activity: **Fig. 4c**).

**Fig. 4:**
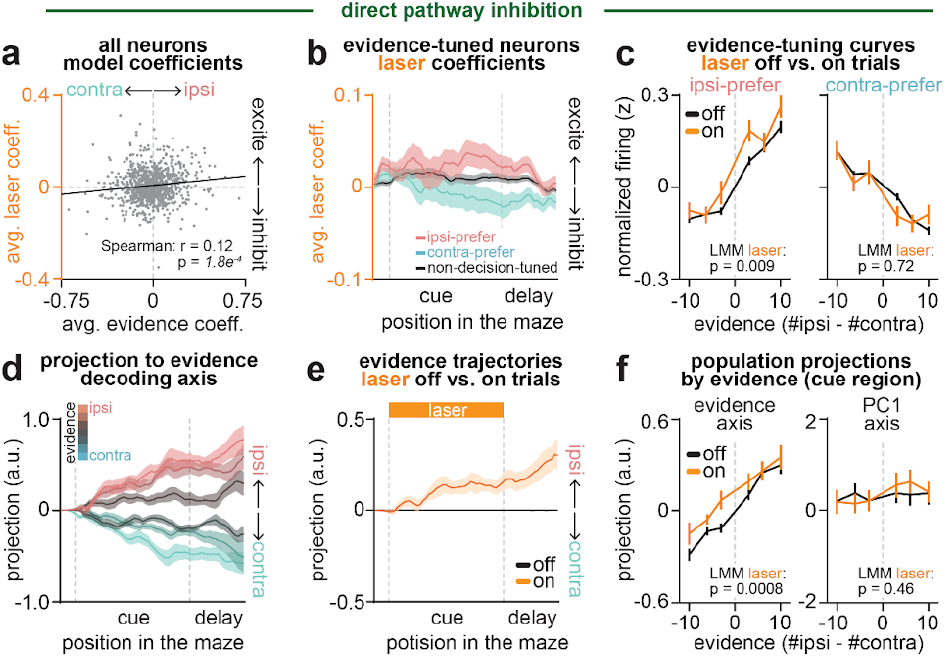
Direct pathway inhibition shifts ACC activity towards ipsilateral-evidence representations. **a**, Average evidence vs. laser coefficients across cue and delay positions in the task for all ACC neurons in direct pathway mice (grey dots; n = 1449 neurons from 14 sessions). Black line: Spearman rank correlation; r = 0.12; p < 0.001. **b**, Averaged laser coefficients at each position in the task for all ACC neurons that showed significant tuning for ipsilateral evidence (salmon; n = 111), contralateral evidence (aqua; n = 130), or neurons without significant tuning to evidence or choice (black, n = 750). **c**, Evidence tuning curves for all ACC neurons with significant evidence-tuning, parsed by ipsilateral-(*left*) or contralateral (*right*) preference. For each neuron, z-scored activity was averaged in all cue and delay positions across trials binned by sensory evidence level, separately for laser off (black) or on (orange) trials. Statistics reflect significance of a linear mixed-effects model (ipsilateral: p_laser_<0.01; contralateral: p_laser_=0.72). **d**, Cross-validated projection of ACC population activity onto a one-dimensional evidence decoding axis at each position in the maze (n = 12 sessions with more than 50 neurons), and then averaged across laser off trials binned by their sensory evidence level. **e**, Same as **d**, but for the averaged evidence-axis projection across all held-out laser off (black) or on (orange) trials. Positive (or negative) values (a.u., arbitrary unit) indicate greater ipsilateral (or contralateral) evidence representation. **f**, Similar to **c**, but evidence tuning curves from population activity projected onto the evidence-axis (*left*) or first principle component (PC1)-axis (*right*). Tuning curves were constructed by averaging projected population activity across positions in the cue region and trials binned by sensory evidence level (#ipsilateral - #contralateral cues), separately for laser off (black) or on (orange) trials. Statistics reflect significance of a linear mixed-effects model; evidence axis: p_laser_<0.001; PC1 axis: p_laser_=0.46. Data are presented as mean ± s.e.m.

Consistent with these single neuron effects, at the population level, direct pathway inhibition produced significant shifts in activity projected onto the evidence decoding axis toward greater ipsilateral evidence representations (**Fig. 4d-f**; **Extended Data Fig. 6h**). This is consistent with the ipsilateral behavioral bias elicited by this manipulation (**Fig. 1e**; **Extended Data Fig. 6l**), and opposite to the contralateral neural and behavioral shifts induced by indirect pathway inhibition (**Fig. 1e**; **Fig. 3g-i**). As with the indirect pathway, the effect of direct pathway inhibition appeared specific to the decision axis, as the manipulation elicited no significant shifts in population activity along the first principal component dimension (**Fig. 4f**, *right*; **Extended Data Fig. 6i-k**).

Together, these results indicate that indirect and direct pathways of the DMS exert opponent influence over evidence representations in ACC, with the indirect pathway shifting representations towards ipsilateral evidence and the direct pathway towards contralateral evidence.

### Striatal pathways have stronger influence over evidence-tuned than choice-tuned or untuned ACC neurons

We next examined how pathway-specific inhibition impacted ACC neurons with binary tuning to the behavioral choice (**Fig. 2a-c**, *bottom row*). Compared to evidence-tuned neurons (**Fig. 3d**; **Fig. 4a**), inhibiting either pathway had weaker effects on activity in choice-tuned neurons. Indirect pathway inhibition elicited a small, but significant, excitation of choice-tuned neurons with contralateral preference (**Fig. 5a-b**, *top*), while direct pathway inhibition had no significant effects on choice-tuned neurons with either preference (**Fig. 5a-b**, *bottom*).

**Figure 5:**
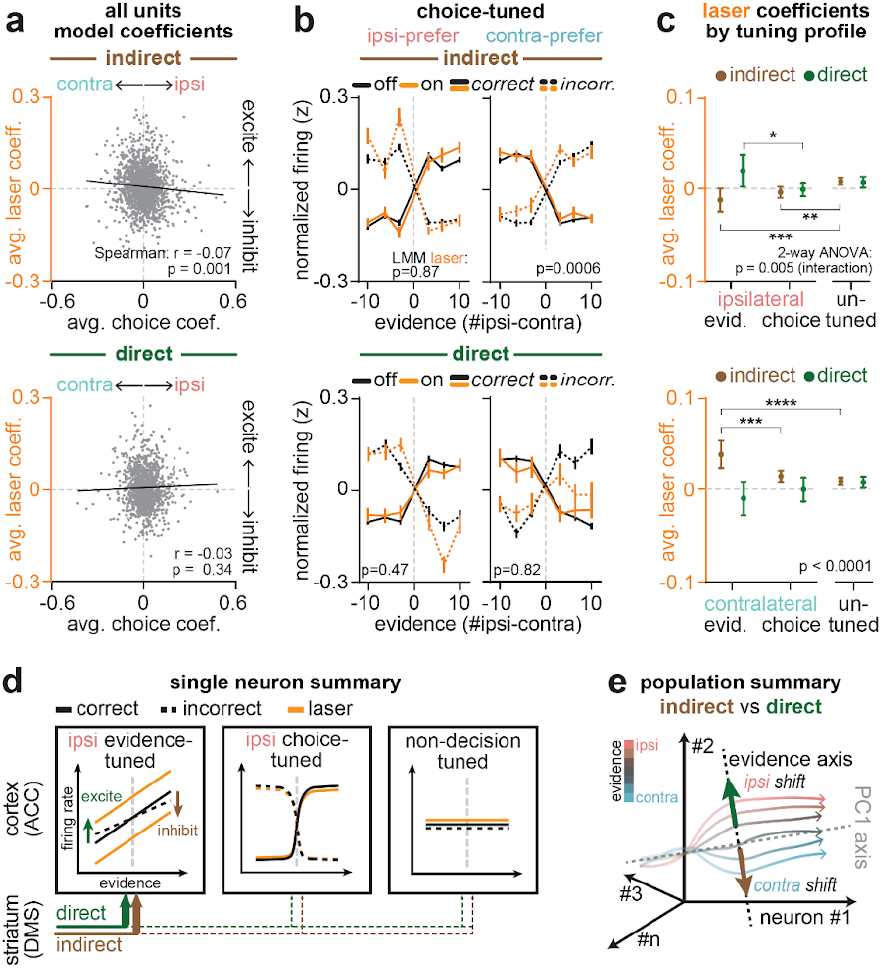
Striatal pathways have stronger influence over evidence-tuned than choice-tuned or untuned ACC neurons. **a**, Average choice vs. laser coefficients across cue and delay positions in the task for all ACC neurons recorded with indirect (*top*; grey dots; n = 2030 neurons from 12 sessions) or direct (*bottom*; grey dots; n = 1449 neurons from 14 sessions) pathway inhibition. Black line: Spearman rank correlation; indirect: r = −0.07, p < 0.01; direct: r=-0.05, p=0.11. **b**, Evidence tuning curves for all ipsilateral-(*left*) and contralateral-preferring (*right*) choice-tuned ACC neurons with indirect (*top*) or direct (*bottom*) pathway inhibition. For each neuron, z-scored activity was averaged in all cue and delay positions and across trials binned by sensory evidence level, separately for laser off (black) and on (orange) trials (indirect: n = 401 ipsilateral-choice, n = 415 contralateral-choice neurons; direct: n = 262 ipsilateral-choice, n = 196 contralateral-choice neurons). Statistics reflect significance of a linear mixed-effects model (indirect, ipsilateral: p_laser_=0.87; indirect, contralateral: p_laser_<0.001; direct, ipsilateral: p_laser_=0.47; direct, contralateral: p_laser_=0.82). Data are presented as mean ± s.e.m. **c**, Average laser coefficient across the cue and delay regions for all ACC neurons with significant tuning to ipsilateral-evidence (indirect: n = 136, direct: n = 111) or ipsilateral-choice (indirect: n = 401, direct: n = 262), or neurons without significant tuning to either choice or evidence (un-tuned; indirect: n = 938; direct: n = 750) *(top*). Brown and green bars reflect ACC neurons from mice receiving indirect or direct pathway inhibition, respectively. *Bottom*: Same as *top*, but for ACC neurons with significant tuning to contralateral-evidence (indirect: n = 140, direct: n = 130) or contralateral-choice (indirect: n = 415, direct: n = 196), or neurons without significant tuning to either choice or evidence (un-tuned; same population as in *top*). P-values reflect significance of a two-way ANOVA for an interaction between tuning (choice-, evidence-, or un-tuned) and pathway inhibition (direct or indirect). Ipsilateral-preferring neurons: p_interaction_<0.01 (F_2,2592_=5.289), p_tuning_<0.05 (F_2,2592_=3.096), p_pathway_<0.01 (F_1,2592_=9.934); contralateral-preferring neurons: p_interaction_<0.0001 (F_2,2563_=10.46), p_tuning_=0.42 (F_2,2563_=0.8630), p_pathway_ <0.0001 (F_1,2563_=26.14). Asterisks denote significance of post hoc unpaired t-tests between groups. Data are presented as a mean ± 95% confidence interval. **d**, Schematic summary of the effects of indirect (brown) and direct (green) pathway inhibition on ACC neurons based on single neuron tuning properties. **e**, Schematic summary of the effects of indirect (brown arrow) or direct (green arrow) pathway inhibition on population activity projected onto the evidence-axis (black dotted line) versus PC1-axis (gray dotted line). *p<0.05, ***p<0.001 and ****p<0.0001.

Indeed, a direct comparison of laser coefficients across ACC subpopulations revealed a large and opposing influence of each pathway on evidence-tuned neurons, which was significantly weaker in choice-tuned neurons (**Fig. 5c**). Weak and non-opposing effects of pathway inhibition were also observed in ACC neurons without tuning to choice or evidence (“un-tuned”, **Fig. 5c**; **Extended Data Fig. 7a**).

The weak effects of the pathways on choice-tuned neurons may seem surprising, given the large changes in the animal’s choice induced by the manipulation (**Fig. 1e**). However, our analyses of the choice-tuned neurons account for choice, either through the encoding model, or by separating trials based on whether they were correct or incorrect. Therefore, this indicates that, while inhibition of each pathway biases choices (**Fig. 1e**), the pathways have little influence on the activity of choice-tuned neurons beyond that explained by the changing choice.

To further clarify how the pathways could affect evidence-tuned neurons and behavior but not the choice-tuned neurons, we simulated models that instantiate different hypothesized relationships between evidence- and choice-tuned neurons. Our data is consistent with a model in which the pathways are primarily having their effect on the evidence-tuned subpopulation, and the choice-tuned neurons are downstream of evidence-tuned neurons, and potentially also downstream of other decision-making processes (**Extended Data Fig. 8**).

Taken together, the specificity of pathway influence on evidence coding in ACC is consistent with an updated classic model (**Fig. 1a**), in which the direct and indirect pathways do not promote and suppress overall cortical activity, but instead selectively and oppositely shift cortical activity along behaviorally-relevant dimensions in each loop. Within the DMS-ACC loop, our results indicate this dimension involves the representation of accumulated visual evidence to guide decisions (**Fig. 5d,e**).

### Effects of pathway inhibition on striatal neurons resemble those in ACC

We also examined how pathway inhibition affected dorsal striatal neurons. Similar to what we observed in ACC (**Fig. 1g-h**), pathway inhibition elicited heterogeneous responses across the entire population of striatal neurons (example neurons: **Extended Data Fig. 2b**; all neurons: **Extended Data Fig. 2d-e**; proportion modulated: **Extended Data Fig. 2f**). Given that we were directly inhibiting DMS neurons, excitatory responses were likely a product of synaptic effects, due to local striatal connections^48–51^ and/or nonlocal inputs from ACC and elsewhere^36,52,53^.

When we instead only considered evidence- or choice-tuned neurons (**Extended Data Fig. 9a-b**), we also observed similarities to the effects of pathway inhibition in ACC (**Fig. 3-5**). In particular, evidence-tuned striatal neurons with ipsilateral preference were excited by indirect pathway inhibition, while those with contralateral preference were excited by direct pathway inhibition. As in ACC, these effects produced opponent shifts to striatal population activity projected to the evidence axis but not the first principal component axis (**Extended Data Fig. 9c**). Given that our inhibition did not target DMS neurons based on their tuning properties, the specificity of pathway inhibition effects on dorsal striatum is surprising, and suggests that the functional tuning of ACC and dorsal striatal neurons may be tightly coupled and interdependent^54,55^.

## Discussion

Here, we tested the classic model of cortico-basal ganglia interactions (**Fig. 1a**) in the context of an evidence accumulation task. We find precise and opposite effects of DMS pathways on the representation of accumulated evidence in ACC (**Fig. 3-4**), with comparatively weak and heterogeneous effects on overall activity (**Fig. 1**), choice-tuned neurons (**Fig. 5**), and the first principal component of the population activity (**Fig. 3-4**).

The precise control that DMS pathways exert on the representation of accumulated evidence in ACC could provide a mechanistic explanation for the selective effects of inhibiting each pathway on evidence-based decisions, without affecting movement or sensory-guided decisions in the absence of evidence accumulation^9,32^. These effects on cortical activity are likely mediated through the basal ganglia-recipient thalamus, which have been shown to have preferential influence over task-encoding cortical neurons^56–60^.

The question then arises of *why* the DMS pathways selectively affect accumulated evidence representations in the cortex. DMS may play a critical role in creating visual evidence representations in the frontal cortex during initial task-learning. Indeed, DMS is the striatal subregion that receives direct inputs from the visual cortex^61,36,35,37^, and learning selectively strengthens visually-evoked activity in DMS^62^. In addition, dopamine projections from the midbrain to DMS have strong pre-training visual responses^63,64^, which have been implicated in the learning of another visual decision-making task^63^.

More broadly, DMS may be important for learning to orient to goals, including not only visual but also auditory^32^ and internally-generated goals^65–75^. Traditionally, with overtraining, DMS-dependent learning is thought to consolidate to dorsolateral striatum (DLS) as behavior becomes habitual^76,77,27,78–80^. While this likely explains why sensory-guided behaviors do not require DMS after training^9,81^, this consolidation process does not appear to occur for evidence accumulation tasks, which remain DMS-dependent even after training^32,9,54^. The fact that such tasks are hard for animals to learn might relate to why they are more difficult to consolidate out of DMS.

Taken together, our results lead to an updated classic model, in which the two basal ganglia pathways oppositely shift behavior and cortical coding for the *relevant* behavioral variable, which may be accumulated evidence in DMS. This may reflect a more general principle, in which the two pathways within other parallel cortico-basal ganglia circuits oppositely control the cortical encoding of different behavioral variables, such as threat learning in the tail of the striatum^82^, Pavlovian associations in the nucleus accumbens^83^, and finer aspects of sensorimotor control in the dorsolateral striatum^84–86^.

**Extended Data Fig. 1:**
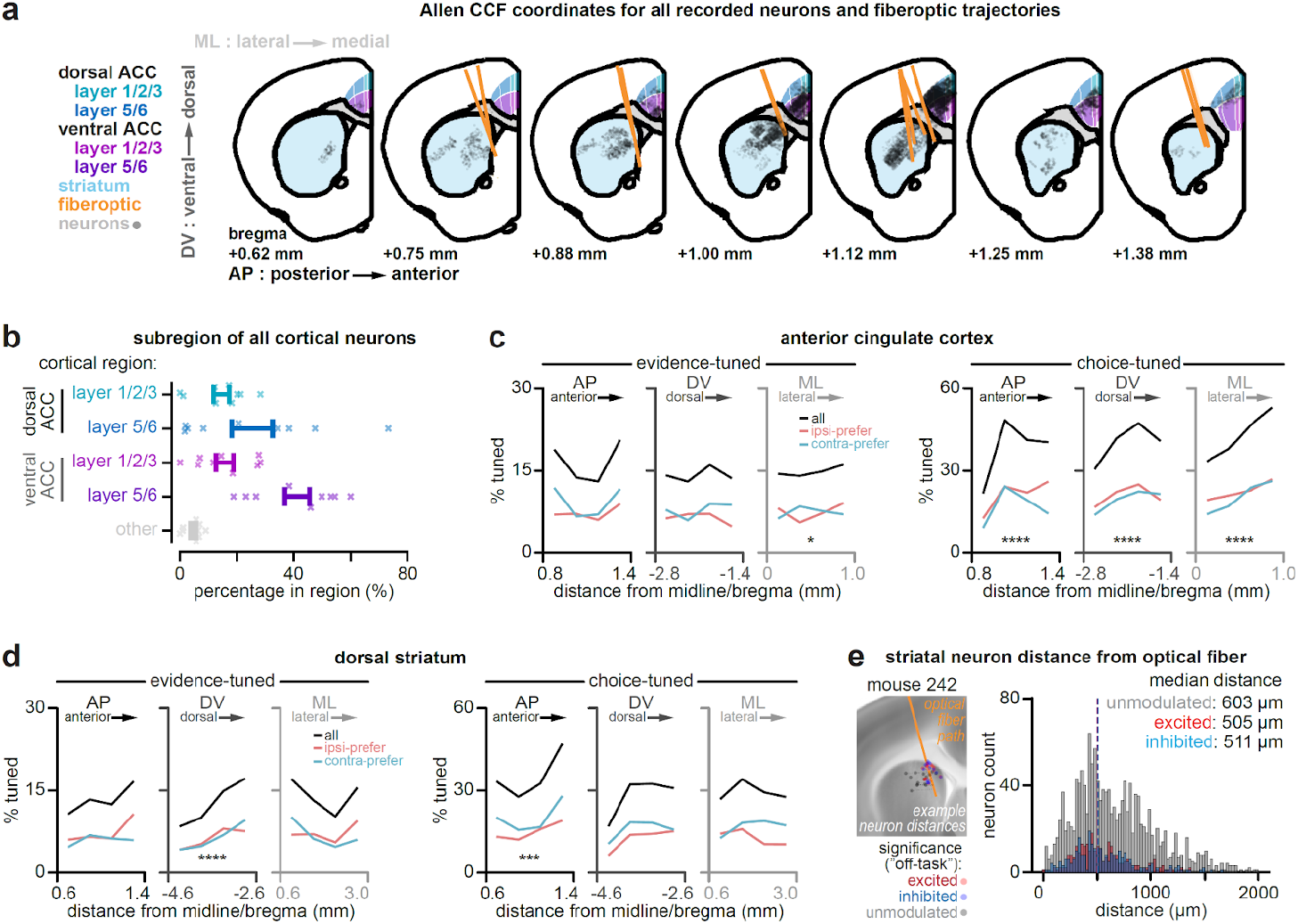
Registration of optical fiber and electrode tracks to the Allen Common Coordinate Framework. **a**, Registration to the Allen Common Coordinate Framework (CCF) of all optical fibers (orange line) and neurons (grey transparent dots) from all mice (n = 14). Dorsal ACC layers (dark blue), ventral ACC (purple), and striatum (light blue). **b**, Proportion of cortical neurons localized to Layer 1/2/3 (light blue) or Layer 5/6 of dorsal ACC (dark blue), Layer 1/2/3 (magenta) or Layer 5/6 of ventral ACC (purple), or other cortical areas (grey). **c**, Proportion of ACC neurons significantly tuned to evidence (*three left columns*) or choice (*three right columns*) based on their AP, DV, or ML CCF coordinates. Proportions are displayed for all significantly tuned neurons (black lines) or separately for neurons with significant ipsilateral-(salmon) or contralateral-(aqua) preference. For each anatomical axis, the full range of observed coordinates were divided into four equally spaced spatial bins. Statistics reflect significance of a linear mixed-effects model to predict tuning (1, tuned; 0, non-tuned) based on each neuron’s CCF coordinates (evidence-tuned neurons: p_AP_=0.19, p_DV_=0.31, p_ML_<0.05; choice-tuned neurons: p_AP_<0.0001, p_DV_<0.0001, p_ML_<0.0001). **d**, Same as **c**, but for dorsal striatal neurons (evidence-tuned: p_AP_=0.44, p_DV_<0.0001, p_ML_=0.94; choice-tuned: p_AP_<0.001, p_DV_=0.36, p_ML_=0.62). **e**, Example (*left*) of a registered optical fiber path (orange line), and the location of example neurons colored by their response to pathway inhibition (red: excitation; blue: inhibition; grey: unmodulated). Dotted lines reflect the shortest estimated distance from each neuron to the optical fiber path. Histogram (*right*) of estimated distances to the optical fiber for all striatal neurons, displayed separately for excited (red), inhibited (blue), or unmodulated (grey) neurons.

**Extended Data Fig. 2:**
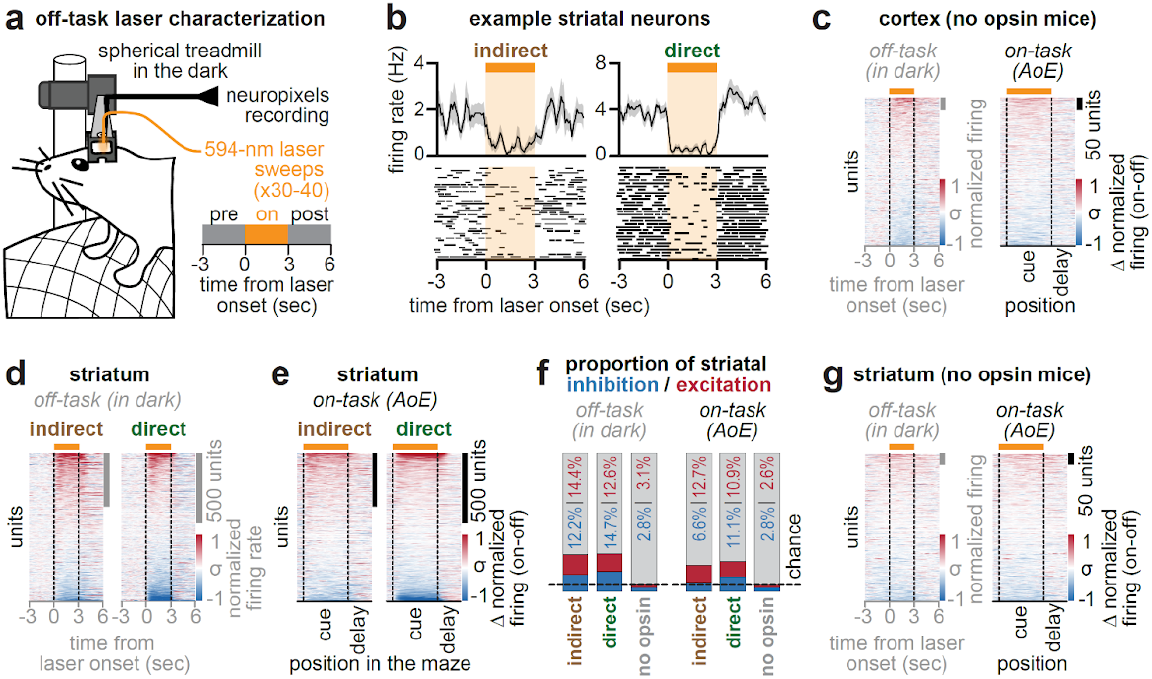
Characterization of pathway-specific inhibition of the striatum. **a**, Schematic of recording and laser delivery (30-40 sweeps, 3 s on, 7 s ITI) as head-fixed mice ran in the dark (“off task”). Laser sweeps occurred after mice completed accumulation-of-evidence task sessions. **b**, Trial-averaged firing rates (*top row*) and rasters (*bottom row*) from two example striatal neurons that showed reversible inhibition by indirect (*left column*) or direct (*right column*) pathway inhibition in the “off task” setting. **c**, Trial-averaged activity for all ACC neurons recorded in the “off task” (*left*) or “on task” (*right*) conditions in mice receiving laser illumination in the absence of opsin expression (n = 509 neurons from 2 mice and 5 sessions). Activity is z-score normalized and sorted by the magnitude of laser modulation in each task condition. **d**, Same as **c**, *left*, but for all striatal neurons recorded from mice receiving indirect (*left*: n = 1135 neurons from 19 sessions) or direct (*right*; n = 836 neurons from 13 sessions) pathway inhibition during the off-task condition. **e**, Same as **c**, *right*, but for all striatal neurons recorded from mice receiving indirect (*left*) or direct (*right*) pathway inhibition in the “on task” condition. **f**, Proportion of significantly excited (red) or inhibited (blue) striatal neurons as a function of task context *(left*: off-task; *right*: on-task) and inhibition group (indirect, direct, or no opsin control). **g**, Same as **c**, but for all striatal neurons (n = 541 neurons from 5 sessions) recorded from no opsin control mice (n = 2 mice).

**Extended Data Fig. 3:**
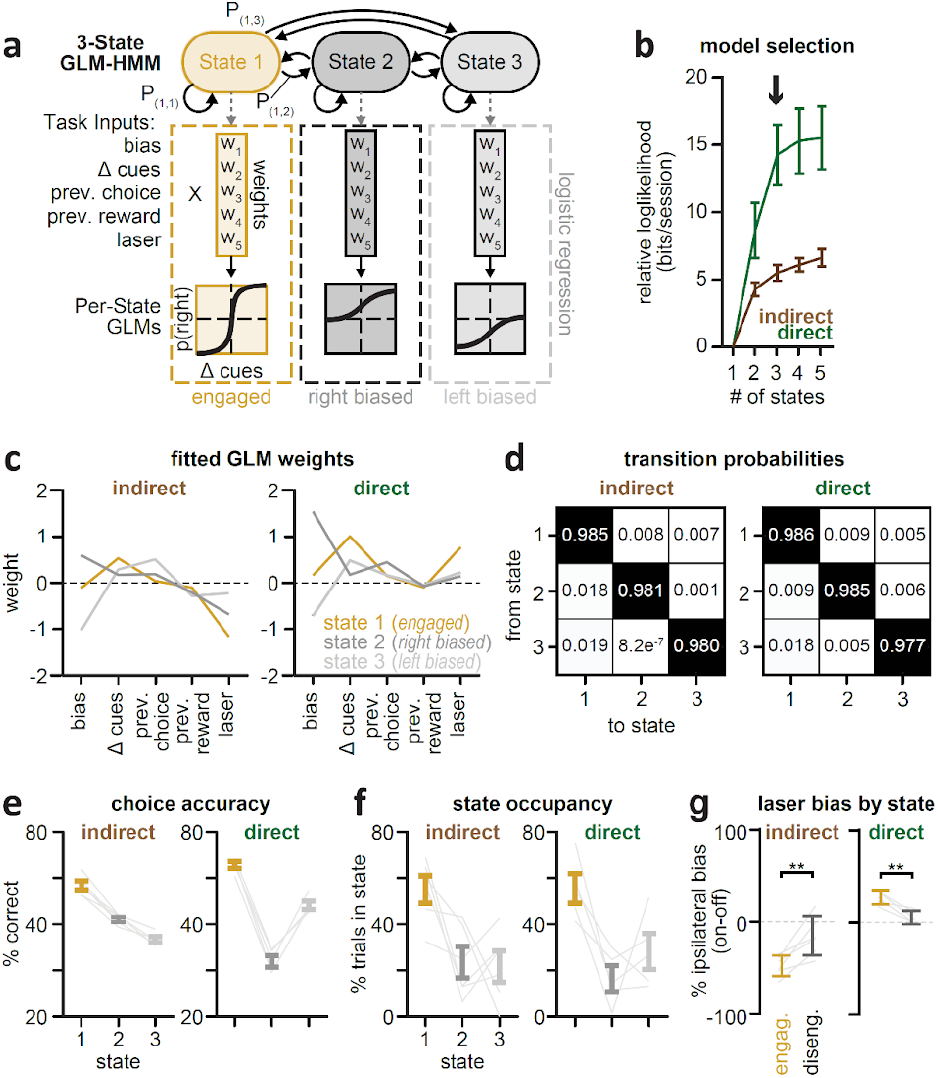
Characterization of GLM-HMM fits to behavioral data. **a**, Schematic of hidden Markov Model with Generalized Linear Model observations (GLM-HMM). The model has latent states with fixed probabilities of transitioning between them, and each state is associated with unique GLM weights that map task inputs (Δ cues, previous choice, previous rewarded choice, laser, and a bias term) onto the probability of a choice (right or left). **b**, Comparison of the cross-validated log-likelihood of the data using GLM-HMMs with different numbers of states, separately for mice receiving indirect (brown) or direct pathway (green) inhibition. All values are relative to the log-likelihood of the 1-state GLM-HMM, and are expressed as bits per session. Solid curves denote mean ± s.e.m. across five different held-out test sets. **c**, Fitted GLM weights for the 3-state GLM-HMM for mice receiving indirect (*left*) or direct (*right*) pathway inhibition. The magnitude of the weight represents the relative importance of that covariate in predicting choice, whereas the sign of the weight indicates the choice direction (positive, rightward; negative, leftward). **d**, Matrix of learned transition probabilities for transitioning from each state to all others on a given trial, separately for mice receiving indirect (*left*) or direct (*right*) pathway inhibition. **e**, Overall choice accuracy (% correct) in each state. **f**, State occupancy, or the percentage of total trials in which each state contained the highest posterior probability. **g**, The difference (on-off) in ipsilateral choice bias (% correct on ipsilateral-contralateral trials) in each state. State 2 (right-biased) and State 3 (left-biased) were pooled into a single “disengaged” state to increase the number of trials for comparison to State 1 (“engaged”). Error bars denote mean ± s.e.m in each mouse; grey lines indicate mean for individual mice; significance was tested using a two-tailed paired t-test (indirect: p<0.01, t(8)=4.7; direct: p<0.01, t(10)=-3.3).

**Extended Data Fig. 4:**
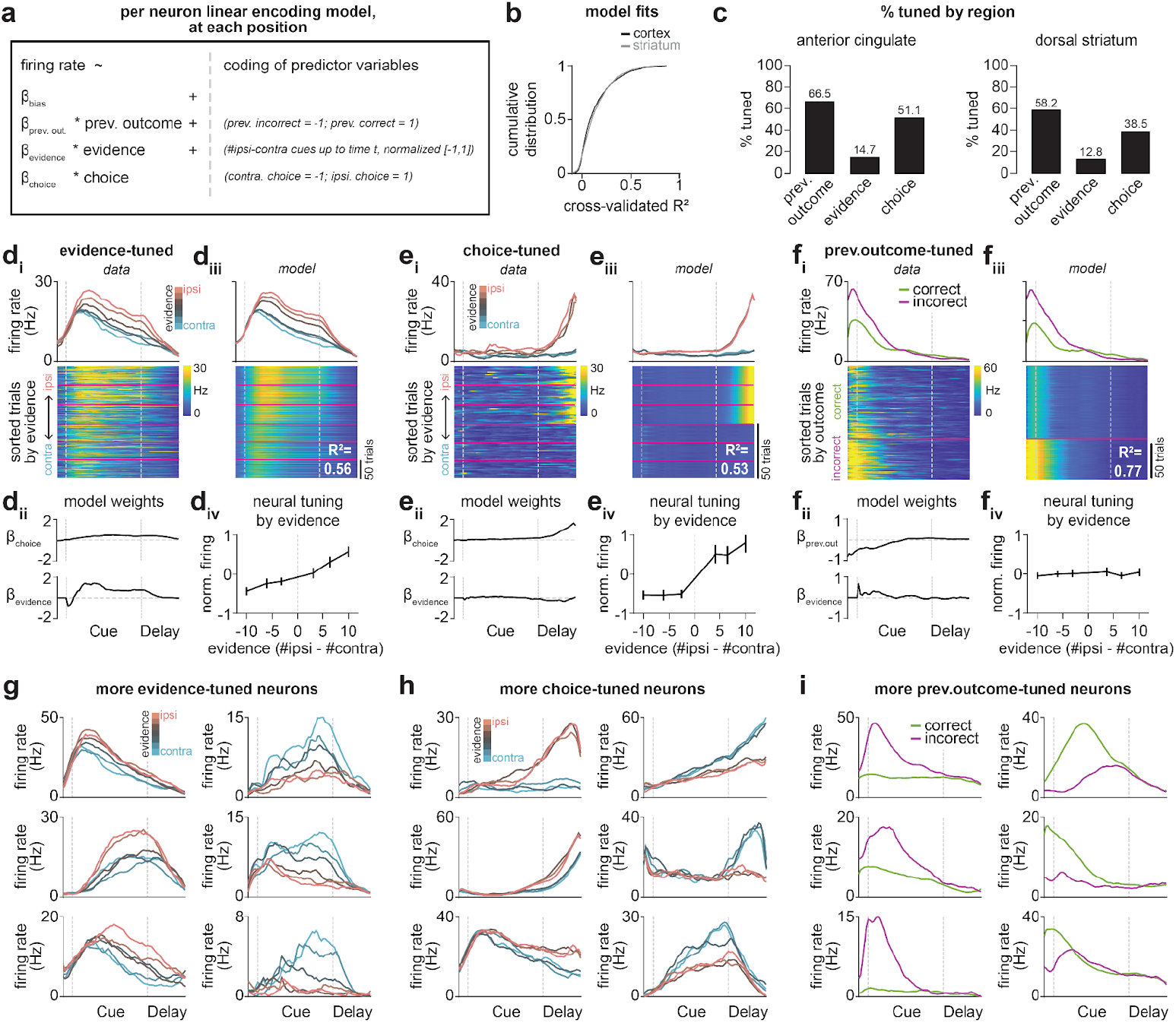
Linear encoding models for identifying task-relevant ACC and dorsal striatal neurons in the accumulation-of-evidence task. **a**, Linear encoding model used to predict the trial-by-trial activity of each neuron at each position based on task variables. All correct/incorrect laser off trials were used to fit the model. **b**, Cumulative distribution of cross-validated coefficient of determination (r^2^) between neural activity and model-predicted neural activity for all ACC (*black* line; n = 3147 neurons from 27 sessions) and dorsal striatal (*grey* line; n = 2136 neurons from 23 sessions) neurons. **c**, Proportion of all ACC (*left*) and striatal (*right*) neurons that showed significant tuning to each task variable. Note that single neurons can be tuned to multiple task variables. **d**, An example neuron with significant tuning to evidence. d*i*, Position-binned firing rate (Hz) in the task (*top*), averaged across trials binned by their final sensory evidence level (#ipsilateral-contralateral cues). Heatmap of trial-by-trial position-binned activity (Hz) for this neuron (*bottom*), with trials sorted by sensory evidence level. Vertical white-dotted lines indicate the start of cue and delay regions in the task. Red horizontal lines (*bottom*) indicate bins of trials by sensory evidence level (as used in *top*). d*ii*, Model coefficients for choice (*top*) or evidence (*bottom*) across position in the task. d*iii*, Same as d*i*, but for model-predicted neural activity. d*iv*, Evidence tuning curve, constructed by averaging neural activity in the cue and delay regions across trials binned by sensory evidence level (as in d*i*). Error bars denote mean ± s.e.m across trials. **e**, Same as **d**, but for an example neuron with significant tuning to choice, and using correct trials only. **f**, Same as **d**, but for an example neuron with significant tuning to previous outcome. **g**, Position-binned firing rate (Hz) in the task for six example neurons with significant tuning to evidence, averaged across trials binned by their sensory evidence level (#ipsilateral-contralateral cues). **h**, Same as **g**, but for six example neurons with significant tuning to choice. Note that only correct trials were included here. **I**, Same as **g**, but for six example neurons with significant tuning to previous outcome (correct, green; incorrect, purple).

**Extended Data Fig. 5:**
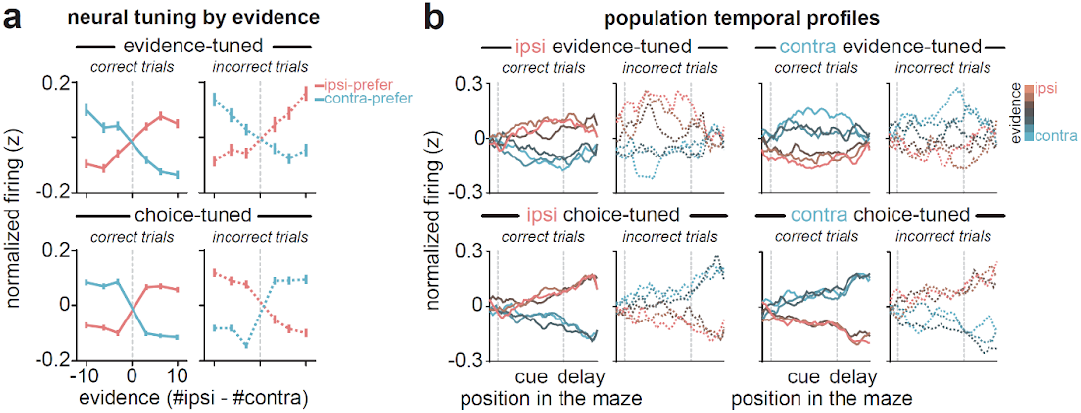
Decision-coding neurons in the dorsal striatum. **a**, Evidence-tuning curves averaged across all evidence-(*top row*; n = 141 ipsilateral- and 133 contralateral-preferring), or choice-tuned striatal neurons (*bottom row*; n = 301 ipsilateral- and 371 contralateral-preferring). Neurons were grouped by their ipsilateral (salmon) or contralateral (aqua) side preference. For each bin of sensory evidence, z-scored activity was averaged across all cue and delay positions in the task, separately for correct (*left*) or incorrect (*right*) trials (n = 2136 total striatal neurons across 23 sessions). Lines and error bars denote mean ± s.e.m. across neurons. **b**, Averaged z-scored firing rate across position in the task for all evidence-(*top row*) or choice-tuned (*bottom row*) striatal neurons, parsed by their ipsilateral (*left two columns*) or contralateral (*right two columns*) preference, and displayed separately for correct (solid lines) and incorrect (dashed lines) trials.

**Extended Data Fig. 6:**
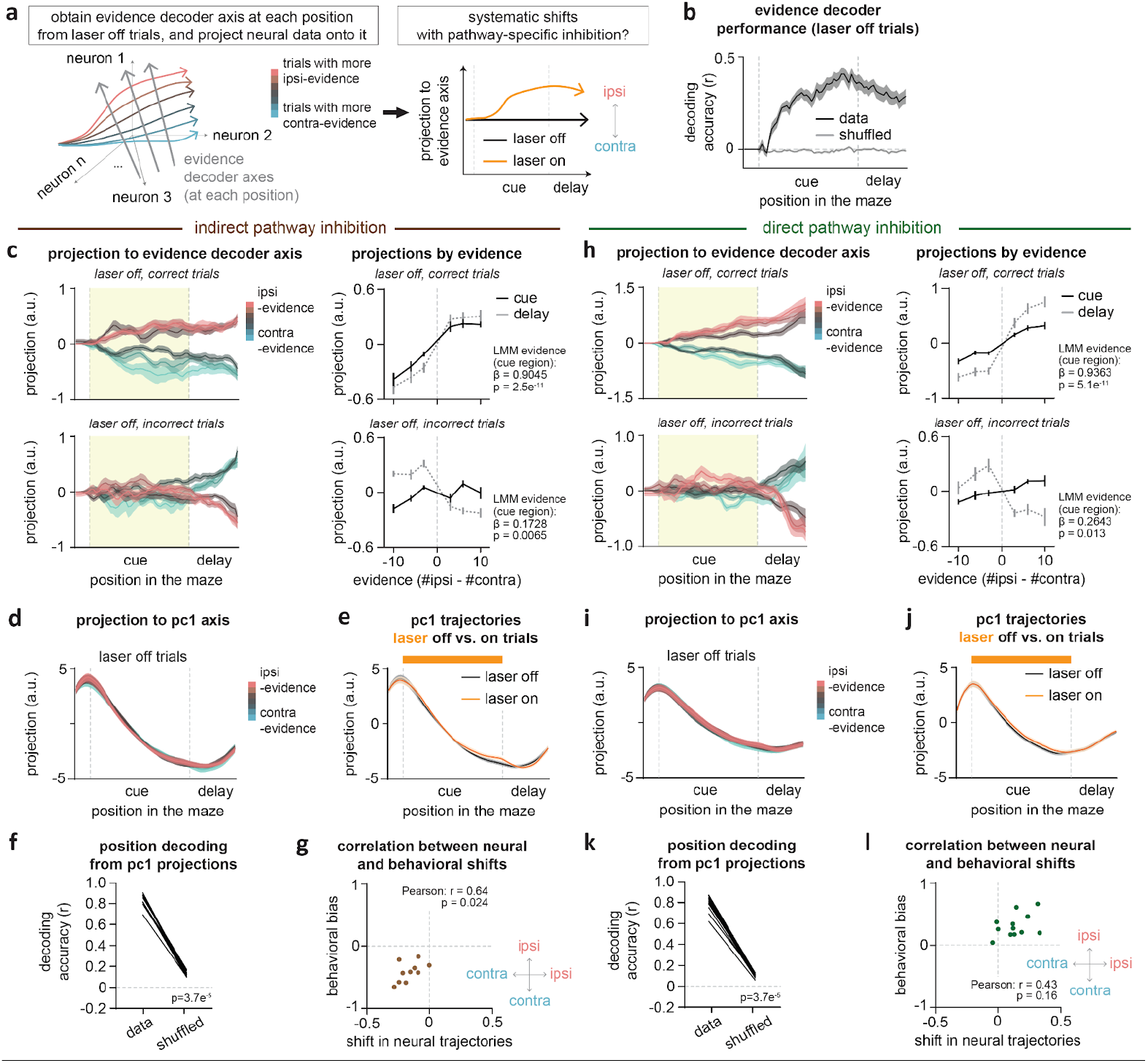
Additional details on the population projections to the evidence decoder or first principal component axis. **a**, *Left*: Schematic of one-dimensional axes in n-dimensional population activity space (with n equal to the number of simultaneously recorded neurons) along which sensory evidence can be maximally decoded at each position in the accumulation of evidence task (grey arrows). Colored lines indicate the firing rate for three neurons across position/time, averaged across trials binned by their sensory evidence level (#ipsilateral - #contralateral cues; ipsilateral, salmon; contralateral, aqua). *Right*: Schematic of average evidence-axis projection across position/time, separately for all laser off (black) or on (orange) trials. An upward (or downward) shift on laser on (orange) trials would indicate a shift towards greater representation of ipsilateral (or contralateral) evidence. **b**, Cross-validated decoding of sensory evidence level (#ipsilateral - #contralateral cues) based on evidence-axis projections from real data (black) or data in which sensory evidence level was randomly shuffled across trials (grey). Decoding accuracy was assessed by calculating Pearson’s correlation coefficient (r^2^) between actual trial-by-trial sensory evidence level and that predicted by evidence-axis projections. Only laser off trials were included for analysis. Error bars denote mean and s.e.m. across sessions (n = 24 sessions pooled across indirect and direct pathway groups). **c**, In mice that received indirect pathway inhibition, cross-validated projection of ACC population activity onto a one-dimensional evidence decoder axis at each position in the task (*left column*), shown separately for all held-out, laser off, correct (*top*) or incorrect (*bottom*) trials. For individual sessions, evidence projections were averaged across trials with varying levels of sensory evidence (#ipsilateral - #contralateral cues). Error bars denote mean and s.e.m. across sessions (n = 12 sessions). *Right column*: Evidence tuning curves obtained by averaging evidence-axis projections in all position bins in the cue (black) or delay (grey) regions across trials binned by their sensory evidence level, separately for all correct (*top*) or incorrect (*bottom*) trials. Note that within the cue region (yellow boxes at left; black lines on right), evidence-axis projections show a preserved positive linear response to evidence levels on both correct (*top right*, black) and incorrect (*bottom right*, black) trials (similar to evidence-tuned neurons in **Fig. 2b**, *top*). In contrast, the response of evidence projections to evidence level during the delay region is more step-like on correct trials (*top right*, grey), and this relationship is inverted on incorrect trials (*bottom right*, grey) (similar to choice-tuned neurons in **Fig. 2b**, *bottom*). A linear mixed-effects model was used to assess tuning curve slope and significance, as reported in each panel. **d**, As in **e**, but for ACC population activity projected onto the first principal component (PC1) axis and averaged across all held-out laser off trials. **e**, As in **d**, but for population activity projected onto the PC1-axis, averaged across all laser off (black) or on (orange) trials. **f**, Decoding accuracy using PC1-projections to predict the position of mice in the maze, based on either real data or data in which position identity was randomly shuffled. Black lines reflect individual sessions. Mann-Whitney U test: p<0.0001, *U* = 4.1281. **g**, Correlation between the laser-induced difference (on-off) in averaged evidence-axis projections and the difference (on-off) in % ipsilateral choice bias (% accuracy on ipsilateral - contralateral trials) from sessions in mice receiving indirect pathway inhibition. Evidence-axis projections were averaged across position bins 100-200 cm. Pearson’s correlation, r = 0.64, p<0.05. **h**, Same as **c**, but for sessions from mice that received direct pathway inhibition (n = 12 sessions). **i**, Same as **d**, but for mice that received direct pathway inhibition. **j**, Same as **e**, but for mice that received direct pathway inhibition. **k**, Same as **f**, but for mice that received direct pathway inhibition. Mann-Whitney U test: p < 0.0001, *U*=4.1281. **l**, Same as **g**, but for mice that received direct pathway inhibition. Pearson’s correlation: r = 0.43, p = 0.16. All data shown as mean ± s.e.m. across sessions unless otherwise stated.

**Extended Data Fig. 7:**
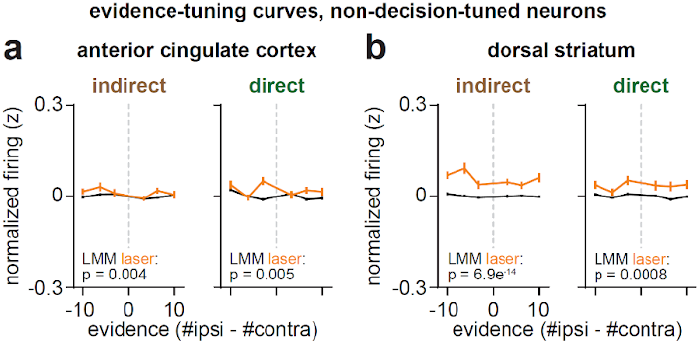
Evidence tuning curves of ACC and dorsal striatal neurons that are neither tuned to evidence nor choice. **a**, Evidence tuning curves for all ACC neurons that did not show significant tuning to evidence or choice, in mice receiving either indirect (*left*; n = 938 neurons from 12 sessions) or direct (*right*; n = 750 neurons from 15 sessions) pathway inhibition. For each neuron, z-scored activity was averaged in all cue and delay positions across trials binned by sensory evidence level (#ipsilateral-#contralateral cues), separately for laser off (black) or on (orange) trials. Statistical significance was assessed with a linear mixed-effects model (indirect: p_laser_<0.01; direct: p_laser_<0.01). **b**, Same as **a**, but for all striatal neurons that did not show significance for evidence or choice (indirect; n = 450 neurons from 19 sessions; p_laser_<0.0001; direct; n = 445 neurons from 13 sessions; p_laser_<0.001). Data are presented as the mean ± s.e.m. (across neurons).

**Extended Data Fig. 8:**
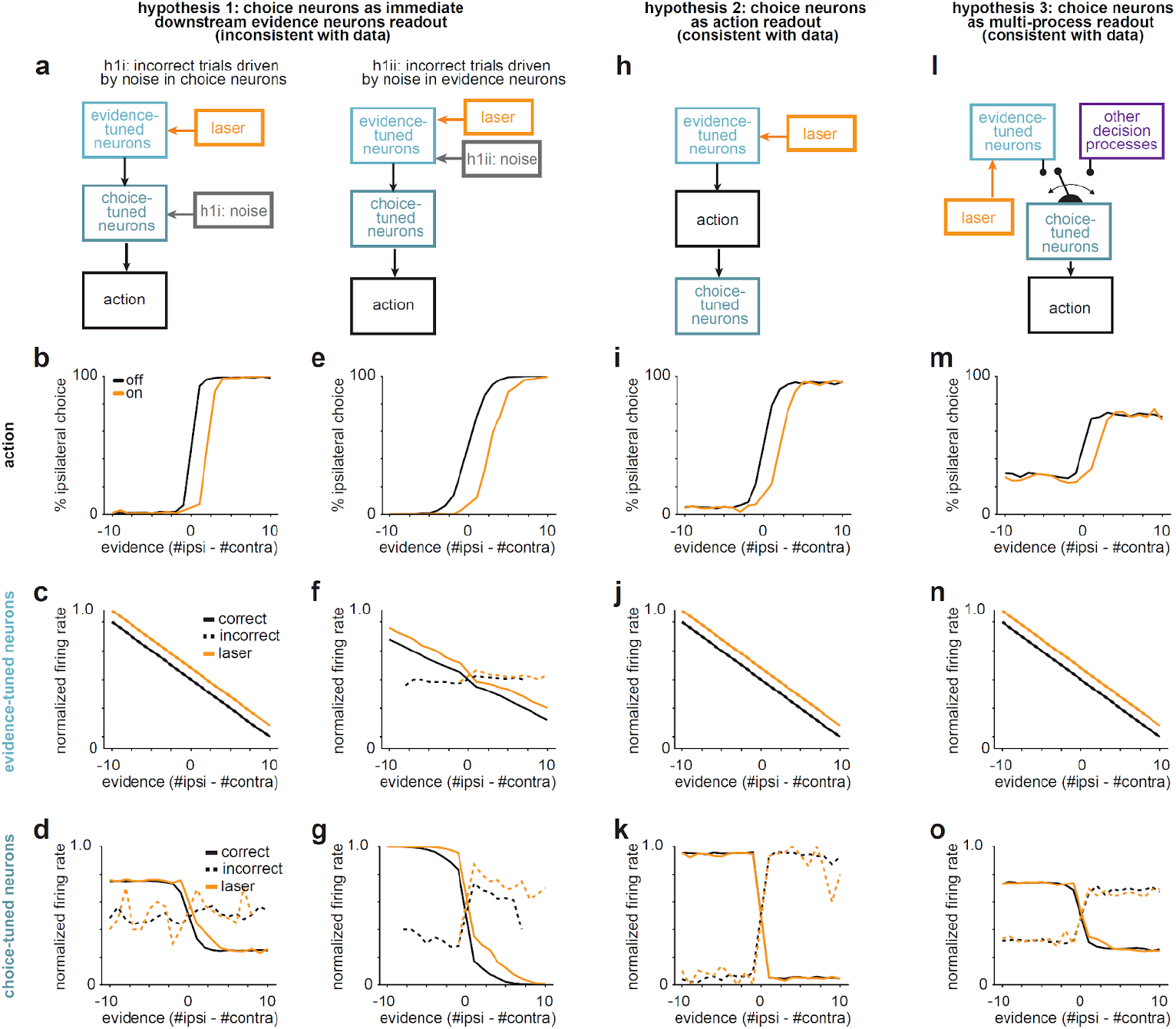
Model simulations of hypothesized relationships between DMS inhibition, evidence- and choice-tuned neurons. **a**, Schematic of model in which laser acts only on the evidence-tuned neurons to shift tuning, and choice-tuned neurons directly readout from evidence-tuned ones with a logistic response pattern. Action is determined by the average of the activity of choice-tuned neurons. Incorrect trials may result from either noise in the choice-tuned neurons’ response to evidence (h1i, *left*) or noise in the evidence-tuned neurons’ response to sensory evidence (h1ii, *right*). **b-c**, Simulations of the model in h1i, showing resulting psychometric curves (b), tuning curves of the noiseless evidence-tuned neurons (c), and tuning curves of the choice-tuned neurons (d). The choice-tuned neurons show responses similar to the data on correct trials with some shifts in tuning near zero but responses on incorrect trials show decreases in amplitude, suggesting the model is incomplete in explaining incorrect trials. **e-g**, Simulations of the model in h1ii showing resulting psychometric curves (e), tuning curves of the evidence-tuned neurons with noise which show patterns inconsistent with the data on incorrect trials, where evidence tuning is flat or has opposite slope (f), and tuning curves of the choice-tuned neurons (g). In both cases, hypothesis 1 fails to capture observed patterns in the data for incorrect trials. This suggests that this mechanistic readout structure is incorrect or that there are patterns of structured noise not accounted for in our simulations. **h**, Schematic of hypothesis 2 in which laser inhibition acts only on the evidence-tuned neurons to shift tuning, and action is determined by a noisy readout of the evidence-tuned neurons. Choice-tuned neurons directly readout from the action. **i-k**, Simulations of hypothesis 2 showing resulting psychometric curves (i), tuning curves of the noiseless evidence-tuned neurons (j), and tuning curves of the choice-tuned neurons (k). These responses are similar to the data on correct and incorrect trials. **l**, Another alternative hypothesis is a hybrid of the two models (hypothesis 3), where choice neurons sometimes respond to evidence neurons and sometimes respond to some alternative decision process (such as, choice history, perseveration, action, etc.). **m-o**, Simulations of hypothesis 3 showing resulting psychometric curves (m), tuning curves of the noiseless evidence-tuned neurons (n), and tuning curves of the choice-tuned neurons (o). These responses are similar to the data on correct and incorrect trials. Both hypothesis 2 and hypothesis 3 show good agreement with the data, suggesting that choice neurons may still be downstream of evidence neurons, either mediated by behavior or alternatively responding to evidence and another decision making process. Critically, in both hypotheses, choice-tuned neurons respond coherently to a common signal, such that amplitude of the response is maintained on incorrect trials. Differences in these hypotheses may be better detected by analyzing whether choice-tuned neurons show a horizontal shift in response to laser for near-zero evidence trials.

**Extended Data Fig. 9:**
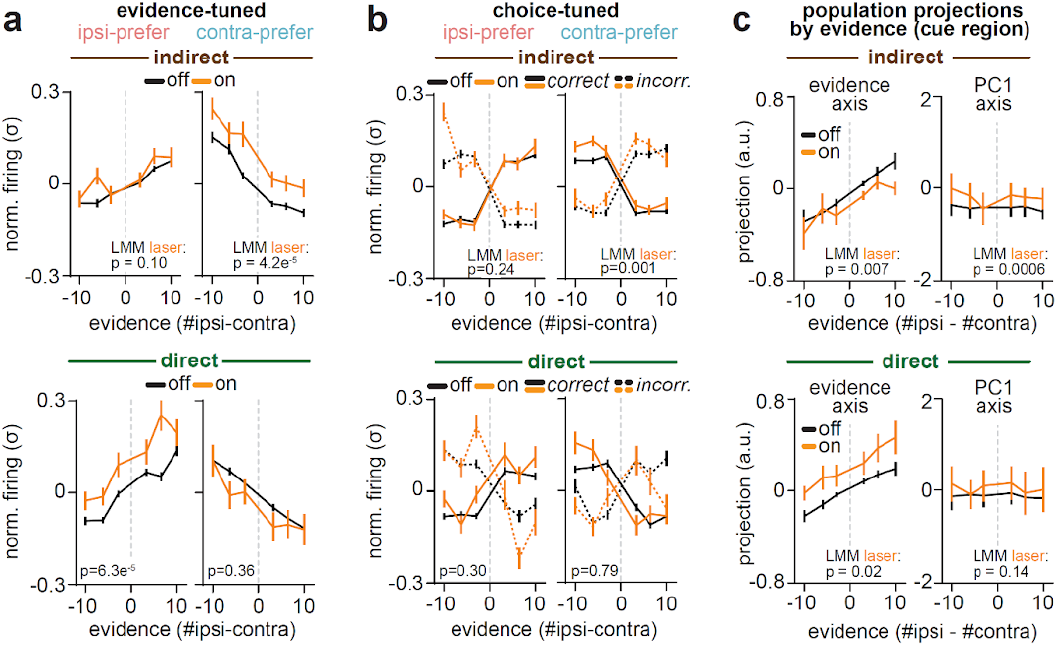
Evidence-tuned neurons in the dorsal striatum show an opponent pattern of modulation in response to pathway-specific inhibition. **a**, In mice that received indirect (*top row*) or direct (*bottom row*) pathway inhibition, evidence tuning curves for all striatal neurons with significant evidence tuning with ipsilateral-(*left column*; indirect: n = 125 neurons; direct: n = 105) or contralateral-preference (*right column*; indirect = 106 neurons; direct: n = 111 neurons). Activity in each neuron was averaged in all cue and delay positions across trials binned by sensory evidence level, separately for laser off (black) or on (orange) trials. **b**, Same as **a**, but for all neurons in striatum with significant choice-tuning, and displayed separately for all correct (solid black and yellow lines) and incorrect (dotted black and yellow lines) trials. Indirect: n = 286 ipsilateral- and 347 contralateral-preferring neurons. Direct: n = 192 ipsilateral- and 160 contralateral-preferring neurons. **c**, Similar to **a**, but for evidence tuning curves obtained from striatal population activity projected onto the evidence-axis (*left column*) or first principal component (PC1)-axis (*right column*) in mice receiving indirect (*top row*) or direct (*bottom row*) pathway inhibition. For all panels, linear mixed-effects models were used to assess statistical significance and p_laser_ is reported in each panel. Data are presented as the mean ± s.e.m. (across neurons in **a** and **b**, or across sessions in **c**).

## Methods

### Animals

We used heterozygous male and female transgenic mice, aged 2-6 months of age, from the following two strains: Drd1-Cre (n = 6, EY262Gsat, MMRRC-UCD) and A2a-Cre (n = 8, KG139Gsat, MMRRC-UCD). Both transgenic lines were backcrossed to a C57BL/6J background (Jackson Laboratory, 000664) and bred in-house. Mice were co-housed with same-sex littermates up until electrode implantation, and subsequently singly-housed in order to avoid damage to the electrode. Mice were maintained on a 12-hr/12-hr light/dark cycle. All surgical procedures and behavioral training occurred in the dark cycle. All procedures were conducted in accordance with National Institute of Health guidelines and were reviewed and approved by the Institutional Animal Care and Use Committee at Princeton University.

### Surgical Procedures

All stereotaxic surgeries were performed under isoflurane anesthesia (3-4% induction, 0.75-1.5% maintenance) after mice received preoperative antibiotics (5 mg/kg Baytril). Mice also received pre- and post-operative analgesia (10 mg/kg Ketofen; 48-hours). Health indicators (i.e. evidence of pain, incision healing, activity, posture) were monitored closely for five postoperative days, and regularly thereafter by experimenters and animal care staff. Upon successful surgical recovery, mice were gradually water-restricted (1-2 mL daily water) and maintained at a minimum of 80% body weight for the duration of the experiment.

#### Viral injection surgery

To inhibit the indirect or direct pathway of the striatum we unilaterally injected the right hemisphere of A2a-Cre or D1R-Cre mice with AAV5-EF1a-DIO-NpHR-eYFP-hGHpA (titer: 1.1-1.8e^13^ genome copies/mL; Princeton Neuroscience Institute Viral Core, or Addgene, 26966). The original viral construct (pAAV-Ef1a-DIO-eNpHR3.0-EYFP)^39^ was a gift from Karl Deisseroth. We used the following coordinates, zeroed from midline/skull surface/bregma, to target the dorsomedial striatum: +1.3 mm ML, −3.2 mm DV, and 0.8 mm and 1.0 mm AP. Virus was delivered at a rate of 100 nl/min and a volume of ∼200 nl at each AP site.

Following viral injection, in the same surgical procedure, the skull was equipped with a titanium headplate (to facilitate head-fixation during virtual reality behavior) and a gold pin (to ground neural signal during subsequent electrode implantation). The gold pin (Newark Electronics) was implanted into the surface of the cerebellum of the left hemisphere, and fixed to the skull with dental cement (Metabond, Parkell S380). The headplate was positioned just dorsal to the skull surface and similarly fixed to the skull with dental cement. A thin layer of dental cement was then applied to the peripheral edges of the skull while leaving the skull surface surrounding bregma exposed. All exposed skull surfaces were then protected with a layer of UV-curing optical adhesive (Norland Optical Adhesive, Batch 61).

Upon post-operative recovery and then gradual water restriction, mice began behavioral shaping procedures in virtual reality (see **Virtual Reality Behavior**, *Shaping*). Upon reaching shaping criteria (∼3-4 weeks), mice were removed from water restriction for a minimum of 24 hours, and then underwent a second surgical procedure for optical fiber and electrode implantation. This gap provided time for viral expression in striatum prior to optogenetic testing, and reduced the duration of electrode implantation and the associated potential for diminished neural signal across time^87,88^.

#### Optical fiber and electrode implantation surgery

To record activity in the ACC and dorsal striatum of the same hemisphere, while inhibiting indirect and direct pathways in this hemisphere, we targeted tapered optical fibers (see **Optogenetics**) to the dorsomedial striatum of the right hemisphere at a 15° angle, while the four shanks of a Neuropixels 2.0 probe (see **Neuropixels Data Acquisition**, *Chronic probe holder*), in line with the coronal plane, were targeted at a −45° angle to traverse the left and then right hemispheres of ACC, and then the right hemisphere of the striatum (**Fig. 1b**).

To achieve this, bregma and the skull surface were first re-exposed by removing UV-curing optical adhesive applied during viral injection surgery. A small craniotomy was made over the right hemisphere to accommodate an optical fiber at the following coordinates: +1.25 mm ML, −3.7 mm DV, and 0.9 mm AP (with 15° angle: +1.92 mm ML, −2.33 mm DV, and 0.9 mm AP). Once the fiberoptic was inserted, the craniotomy was covered with a minimal amount of medical grade petroleum jelly to prevent cement contacting the brain surface, and the fiber was fixed to the skull with dental cement (Metabond, Parkell S380). A large craniotomy was then made over the left hemisphere, at approximately the same AP axis as the fiberoptic, to accommodate the four shanks (750 μm distance between shank 1 and 4) of a Neuropixels 2.0 probe at the following coordinates: +2.5 mm ML, −4.4 mm DV, and 1.0 mm AP (with −45° angle: −1.4 mm ML, −4.7 mm DV, and 1.0 mm AP). The coordinates reflect the brain position of the left-most shank when zeroed at the skull surface of bregma.

Prior to inserting probe shanks into the brain a silver ground wire linking probe ground and reference pads was soldered to the gold pin implanted over the cerebellar surface (see **Surgical Procedures**, *Viral injection surgery*). Once the electrode was inserted, all exposed shanks and brain surfaces were covered in medical grade petroleum jelly and the chronic Neuropixels 2.0 holder was coupled to the skull surface with dental cement and fully encased/protected with fast-curing dentin (Parkell S301).

To limit the visual spread of light when coupling the fiberoptic to the laser, the implant surface was then coated in a final layer of dental resin (Ortho-Jet, Lang Dental 1304) mixed with opaque carbon powder (Sigma-Aldrich, 484164). In order to facilitate histological reconstruction and registration of trajectories to the Allen Common Coordinate framework (**Fig. 1c**; **Extended Data Fig. 1**; see **Histology**), electrode shanks and optical fibers were coated in CellTracker CM-DiI (Invitrogen C7000) prior to implantation. For distances of all recorded striatal neurons from optical fiber trajectories, see **Extended Data Fig. 1e**.

### Virtual Reality Behavior

#### Virtual reality setup

Mice were head-fixed over an 8-inch Styrofoam® ball suspended by compressed air (∼60 p.s.i.) centered within a custom-built toroidal screen spanning a visual field of 270° horizontally and 80° vertically. The setup was enclosed within a custom-designed cabinet built from optical rails (Thorlabs) and lined with sound-attenuating foam sheeting (McMaster-Carr). The virtual reality environment was projected onto the toroidal screen using a DLP projector (Optoma HD141X) with a refresh rate of approximately 60 Hz.

As previously described^6,9^, an optical flow sensor (ADNS-3080 APM2.6), located beneath the ball and connected to an Arduino Due, ran custom code to transform real-world ball rotations into virtual-world movements (https://github.com/sakoay/AccumTowersTools/tree/master/OpticalSensorPackage) within the Matlab-based ViRMEn software engine^89^; http://pni.princeton.edu/pni-software-tools/virmen). The ball and sensor of each VR rig were calibrated such that ball displacements (*dX* and *dY*, where X and Y are parallel to the anterior-posterior and medial-lateral axes of the mouse, respectively) produced translational displacements proportional to ball circumference in the virtual environment of equal distance in each axis. The y-velocity of the mouse was given by 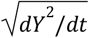, where *dt* was the elapsed time from the previous sampling of the sensor. The virtual view angle of mice was obtained by first calculating the current displacement angle as: ω = *atan*2(− *dX* · *sign*(*dY*), |*dY*|).

Then the rate of change of view angle (*θ*) for each sampling of the sensor was given by:

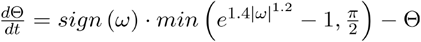

This exponential function was tuned to (i) minimize the influence of small ball displacements and thus stabilize virtual-world trajectories, and (ii) increase the influence of large ball displacements in order to allow sharp turns into the maze arms.

#### Behavioral shaping

Prior to implantation of electrodes for acquiring neural recordings during the evidence accumulation task, mouse behavior was first shaped to perform the task via a series of 9 shaping T-mazes as previously described^6,9^. Criteria for the number of trials completed, percentage of trials with motor errors, overall accuracy, and bias determined progression to a subsequent shaping maze. Daily sessions lasted ∼1-1.5 h, and mice typically completed all shaping mazes in ∼21-28 days.

Prior to the first shaping maze, mice were first handled and habituated to the experimenter, to head-fixation in the VR rig, and to the sucrose reward (10% sucrose in water) for 3-5 days. In all shaping mazes, choosing the correct arm resulted in 4-8 μL of sucrose reward and 3 s ITI, while incorrect choices resulted in no reward and a 12 s ITI. The number and location of visual cues on each trial were drawn randomly according to a spatial poisson process, and were made visible to mice when they were 10 cm from the cue location and remained visible until trial completion in all shaping mazes. Cue positions on the same maze side were also constrained by a 12 cm refractory window to avoid overlapping cues.

A first group of shaping mazes (Maze 1-4) primarily served to shape motor behavior in VR. In these mazes, visual cues were presented on the rewarded T-maze side only, and a prominent visual guide was continuously visible from within the rewarded maze arm itself. Across these mazes, the T-maze stem was gradually extended from an initial length of 60 cm to a full length of 300 cm. In all subsequent shaping mazes the prominent visual guide was removed from the rewarded arm.

In a second group of shaping mazes (Maze 5-7), behavior was shaped to respond to visual cues presented in the T-maze stem, and to withhold a motor turn response during a delay period without visual cues. In Maze 5, visual cues were presented along the entire 300 cm of the T-maze stem up until the choice point, but by Maze 7, visual cues were presented from 0-200 cm only, followed by a delay region (200-300 cm) without cues.

In a final group of shaping mazes (Mazes 8-9), distractor cues on the non-rewarded maze side were introduced with increasing frequency (mean side ratio of rewarded:non-rewarded side cues; maze 8: 8.3:0.7; Maze 9: 8.0:1.6 m^-1^). Upon reaching criterion performance on the final shaping maze (Maze 9), mice progressed to the accumulation-of-evidence task (see below) for a minimum of three sessions before being placed on ad libitum water and subsequently undergoing optical fiber and electrode implantation surgery (see **Surgical Procedures**, *Optical fiber and electrode implantation surgery*).

#### Accumulation-of-evidence task

The task took place in a 330 cm long virtual T-maze with a start region (defined as −30-0 cm), a cue region (0-200 cm), a delay region (200-300 cm), and ending with 90° turns toward left or right choice arms (**Fig. 1d**). In contrast to all shaping mazes, visual cues were presented transiently when mice were 10 cm from their randomly drawn spatial location and disappeared 200 ms later. The mean ratio of reward to non-reward side visual cues presented during the cue region were 8.0 m^-1^:1.6 m^-1^ (∼50% of trials) or 7.3 m^-1^:2.3 m^-1^ (∼32% of trials). A turn toward the maze side with the greater number of cues resulted in 4-8 ul of 10% sucrose reward followed by a 3 s ITI, while a turn toward the T-maze arm with fewer cues resulted in no reward and a 12 s ITI.

To facilitate overall performance, in block-wise fashion we interspersed two additional maze trial types during task sessions. First, each session began with “warm-up” trials, in which a prominent visual-guide was visible in the rewarded arm (see **Virtual Reality Behavior**, *Shaping*, maze 4, ∼7% of trials). Mice progressed to the evidence accumulation maze after ten “warm-up” trials, or until accuracy exceeded 85% correct on them. These trials were excluded from all neural and behavioral analyses. Second, when accuracy fell below 55% over a 40-trial running window within a session, mice transitioned to a maze in which cues were presented only on the rewarded maze side and did not disappear following presentation (see **Virtual Reality Behavior**, *Shaping*, maze 7, ∼11% of trials). These “easy blocks” were limited to ten trials, after which mice returned to the main testing maze regardless of performance, and were included in all analyses to preserve trial history. Optogenetic testing and neural recording sessions typically lasted 1-2 hours and consisted of 200-400 trials.

#### Debiasing algorithm

To discourage side biases, in all shaping and testing mazes we used a previously implemented debiasing algorithm^6,9^. As described in detail elsewhere, this was achieved by changing the underlying probability of drawing a left or right rewarded trial in a pseudo-balanced manner. Briefly, the probability of drawing a right reward trial, P_R_, was given by,

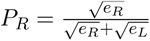

where e_R_ (and e_L_) are the weighted average of the fraction of errors the mouse made in the past 40 right (and left) rewarded trials. To ensure a greater influence of more recent trials we used a half-Gaussian with σ = 20 trials. To discourage the generation of long sequences of only right (or left) rewarded trials, we capped √e_R_ and √e_L_ to be within the range [0.15, 0.85]. Because the empirical fraction of drawn right trials could deviate from P_R_, particularly when the number of trials is low, we applied an additional pseudo-random drawing prescription. Specifically, if the empirical fraction of right trials (calculated using a half-Gaussian window with σ = 60 trials) exceeded P_R_, right trials were drawn with probability 0.5*P_R_, whereas if the empirical fraction was below P_R_, right trials were drawn with probability 0.5*(1+P_R_).

### Optogenetics

To increase the volume of light delivered to brain tissue and provide a larger and more homogenous power density distribution adjacent to electrode recording sites, we used tapered optical fibers fabricated in-house as previously described^9,90^. Briefly, optical fiber tips (Newport F-MBB; 200 μm, 0.37 NA, silica cladding and core) were submerged in hydrofluoric acid (48%, Millipore Sigma), and slowly withdrawn at a rate of 25 μm/min using a gear motor (Newegg, DC 12 V, 1 rpm; run at 5%) to drive a pneumatic micromanipulator (Narishige; MOS-1) coupled to a stereotactic frame (Kopf). The depth of submersion in hydrofluoric acid (∼2.5 mm), the fiber diameter (200 μm), and the rate of withdrawal yielded a tapered conical tip of ∼1.5 mm ± 0.25 mm in length with a ∼3.7° taper angle. Taper quality was assessed by visual inspection of taper length, uniform light emission, and measured power output.

To activate halorhodopsin we utilized a 594 nm diode-pumped laser (Hubner Photonics, Cobalt Mambo 04-01, 100 mW), which coupled to the implanted optical fibers via an optical patch cable fabricated in-house (ThorLabs, 200 μm diameter, 0.37 NA). During behavioral testing, the laser path was gated by a shutter driver (Stanford Research Systems, Model SR474), which was controlled by TTL pulse sent from the PC running the virtual environment. In all experiments, laser delivery (∼2 mW) was restricted to periods when the animal was within the cue region of the maze (0-200 cm), and occurred on a random subset (10-20%) of trials within a session (**Fig. 1d**). The median duration of continuous laser delivery was ∼3.3 s, with a range of ∼2.4 to 6.1 s. Laser was delivered unilaterally to the right hemisphere in all mice, ipsilateral to the recording hemisphere. After completion of behavioral testing, mice received 30-40 laser sweeps while running in the dark (“off-task”). Laser sweeps consisted of 3 s of continuous laser illumination followed by a 7 s laser off inter-trial interval.

We took several steps to limit and control for potential effects of laser illumination to brain tissue, which at high power densities can produce heat capable of influencing neural activity and behavior in the absence of opsin expression^45,46^. First, we used relatively low power levels and durations of laser delivery (∼2 mW; ∼3.3 s), which were both below those reported to have effects on tissue heating and behavior when targeting the dorsal striatum^46^. Second, we used tapered optical fibers with greater light-emitting surface area, which further reduces the local build up of heat at the fiber tip^91,45^. Finally, we directly controlled for potential effects of laser delivery on neural activity by recording activity during behavior in two mice that did not receive viral delivery of halorhodopsin (**Fig. 1h**; **Extended Data Fig. 2c,f,g**).

### Neuropixels Data Acquisition

#### Chronic probe holder

To achieve large-scale and long-term chronic recordings across ACC and dorsal striatum, we utilized test-phase and commercial-release Neuropixels 2.0 electrodes (Imec) containing over 5,000 potential recording sites across 4 shanks with a 0.75 mm by ∼9 mm footprint^88^. To implant electrodes with the possibility for probe recovery and re-use, we utilized a rescaled version of a previously described chronic Neuropixels implant holder^87,92^. The holder design consisted of four discrete parts printed in-house with a Formlabs SLA 3D printer: (i) an internal probe holder containing a stereotaxic adaptor, which coupled to a titanium dovetail fixed to the Neuropixels probe base via a set screw, (ii) a 3-walled external chassis that encased the internal probe holder and coupled to it via machine screws, and (iii) a lid that coupled to the fourth-wall of the external chassis to provide 360° of protective coverage to the internal holder and probe. After assembly, a thin layer of Metabond was applied to the external surface of the chassis, the lid, and their connective points to prevent permanent coupling of internal and external assembly parts and to facilitate subsequent fixation of the chassis to the skull. The dimensions of the probe holder for commercial (or test-phase) probes were approximately: 21 mm (or 24.7 mm) in height, 8 mm (or 12.2 mm) in width, and 12.5 mm (or 11.2 mm) in depth. The maximum weight was ∼1.5 g. External reference and ground pads of all probes were connected via a silver wire during assembly. Design files and instructions for printing and assembling the chronic Neuropixels 2.0 implant are available at https://github.com/agbondy/neuropixels_2.0_implant_assembly.

#### Data acquisition and synchronization

Neural data was acquired at 30 kHz using National Instruments PXl hardware (Pixel-1071, 4-slot chassis; PXLe-6341, X series DAQ; PXle-8381/2, remote control modules; BNC-2110 breakout box) and SpikeGLX software (https://billkarsh.github.io/SpikeGLX) run on a Dell workstation (Precision 3460) with GPU (NVIDIA Quadro, P2200). Imec and NIDAQ data stream clocks were synchronized via a 0.5 s on/off TTL pulse between the Imec SMA connector port and NIDAQ breakout board. Behavioral and neural data were synchronized via TTL pulses sent from the desktop running the virtual environment to the NIDAQ breakout board (a digital TTL pulse at each trial start, and approximately every 16 ms upon each update to the virtual environment).

#### Recording site selection

During behavioral testing we recorded from 384 channels at different spatial-anatomical locations on different days. To select recording sites, we used the activity profile from baseline recordings taken across the most ventral 768 recording sites on each shank for each mouse (5.76 mm in distance from shank tip). From these baseline recordings, we inferred the location in which each shank traversed the following boundaries based on the sparsity or density of neural activity: (i) the corpus callosum of the right hemisphere (defining the border between striatum and cortex), (ii) layer 1 of dorsal cortex on each side of the sagittal midline (defining the border between left and right hemispheres), and (iii) the brain surface (defining the border between the left hemisphere and extracranial space).

For each mouse, we then selected multiple recording banks of 384 channels each, with the goal of minimizing overlap in their recording site locations. Per mouse, we generated non-overlapping recording banks targeting the anterior cingulate cortex only (1-2 recording banks), or the dorsal striatum only (1-2 recording banks). To record from both structures within the same session, we additionally selected 1-2 recording banks in which both cortical and striatal regions were sampled simultaneously. While these cortico-striatal recording banks did not contain overlapping channels with one another, they could sample channel locations included in the region-specific recording banks. For each mouse, each unique recording bank was sampled across 2-6 sessions, separated in time by ∼1-14 days. All recording site locations were verified post hoc using whole-brain clearing, light sheet microscopy, and registration of electrode shank tracks to the Allen Common Coordinate Framework (CCF) based on electrophysiological landmarks^40,41^ (**Fig. 1c**; **Extended Data Fig. 1**; see **Histology**).

#### Spike-sorting

We first used the open-source software package CatGT (https://github.com/billkarsh/CatGT) to (i) remove large artifacts (-*gfix*; minimum absolute amplitude: 0.4 mV; minimum absolute slope: 0.1 mV/sample; noise level defining transient end: 0.02 mV), (ii) apply global common average referencing over all channels (-*glbcar*), (iii) apply a zero-phase high-pass Butterworth filter in the spike frequency range (-*apfilter*; type: butter; order: 12; hi-frequency: 300 Hz; low-frequency: 9000 Hz), and (iv) extract digital TTLs sent from the virtual environment to the NIDAQ data stream (*-xd*). We then used Kilosort 2^93^ (https://github.com/MouseLand/Kilosort) to spike-sort denoised and band-pass filtered data into neuronal clusters. Clusters were manually inspected using phy (https://github.com/cortex-lab/phy) and classified as either single-unit activity, multi-unit activity, or noise based on traditional metrics (e.g. cluster isolation, waveform shape, and inter-spike-interval violations).

#### Removal of duplicate clusters

We took several steps to limit the influence of sampling the same neuron across sessions in our analyses. First, for each mouse we recorded from non-overlapping channel locations across days (see above, *Recording site selection*). Second, when more than one session with the same recording configuration in a mouse was used for analysis, we applied the software package UnitMatch^42^ (https://github.com/EnnyvanBeest/UnitMatch) to identify neurons with matching waveforms across days and only considered such neurons for analysis from a single session.

Specifically, we used the following waveform parameters to generate similarity scores (scaled 0 to 1) between the waveforms of all single-unit and multi-unit activity clusters: (i) the spatial decay of waveform amplitude across recording sites, (ii) the weighted-average waveform, (iii) the weighted-average waveform amplitude, (iv) the average recording site position weighted by the maximum amplitude on each recording site (“spatial centroid”), (v) the trajectory of the spatial centroid −0.2-to−0.5 ms around peak waveform amplitude, (vi) the euclidean distance traveled by the spatial centroid at each time point, and (vii) the travel direction of the spatial centroid at each time point. We then used the average of each parameter’s score to compute a total similarity score, *T*. Based on the distribution of total similarity scores when comparing each cluster to itself – waveforms from the first half of a recording to the second half – versus the distribution of similarity scores for all cross-cluster waveform comparisons, we identified a threshold of *T >* 0.75 to classify true positive waveform matches. Distinct clusters with matching waveforms within a session, which reflects over-splitting during the initial spike-sorting or in post-curation, were then merged into single clusters (∼6.4% of neurons per session). On average, 31.0% of single-unit (and 23.5% of multi-unit) clusters shared matching waveforms across sessions and were considered putatively tracked neurons across days. For all neurons with matching waveforms across sessions, the cluster recorded on the session with the greater number of trials was included for analysis, while all other matching clusters were excluded from single-neuron analyses (**Fig. 1f-h**; **Fig. 3d-f; Fig. 4a-c; Fig. 5a-c; Extended Data Fig. 2, 7 and 9**).

We did not apply this removal procedure in two instances. First, for initial characterization of task-coding neurons on laser off trials (**Fig. 2; Extended Data Fig. 5**), we avoided duplicate clusters altogether by including only one session per unique recording configuration (see *Recording site selection*). For this analysis, we instead selected the session with the greatest number of engaged state trials (see *GLM-HMM*) from each recording configuration. Second, we did not remove duplicate clusters from population decoding analyses (**Fig. 3g-i; Fig. 4d-f; Extended Data Fig. 6**), since in this case the analysis was performed per session, rather than per neuron.

### Behavioral Analyses

#### General performance

Accuracy was defined as the percentage of trials in which mice chose the maze arm corresponding to the side having the greater number of sensory cues (**Extended Data Fig. 3e**). Laser-induced choice bias (**Extended Data Fig. 3g**; **Extended Data Fig. 6g,l**) was measured by first calculating the difference in choice accuracy on trials where sensory evidence indicated an ipsilateral reward versus a contralateral reward on laser on and off trials separately (% correct, ipsilateral–contralateral; positive values indicate greater ipsilateral bias), and then subtracting these two values (% ipsilateral laser bias, on - off; positive values indicate laser induced ipsilateral choice bias). Note that, given the probabilistic delivery of sensory cues, a small number of trials in each session (typically <5/session) contained an equal number of ipsilateral and contralateral cues, and the rewarded arm was randomly assigned. These trials were excluded from all analyses.

#### Psychometric behavior and binning of sensory evidence levels

Psychometric behavior (**Fig. 1e**) was assessed by first defining sensory evidence and choice as either ipsilateral or contralateral relative to the hemisphere of laser delivery, and binning trials based on the total number of #ipsilateral - #contralateral cues presented on each trial (Δ cues). For each session, we grouped all trials (including correct/incorrect, laser on/off) with greater ipsilateral (Δ cues > 0) or contralateral (Δ cues < 0) evidence into three bins with an equivalent number of trials in each group. Each trial grouping thus reflects low, intermediate, and high levels of ipsilateral, or contralateral, sensory evidence. Psychometric curves (**Fig. 1e**) were then calculated as the percentage of ipsilateral choices in each bin for each mouse, separately for all laser on and off trials.

Note that we used the same binning procedure for sensory evidence levels in neural analyses (**Fig. 2-5; Extended Data Fig. 4-7,9**; see **Neural Analyses**, *Evidence tuning curves*). For all behavioral or neural analyses that focused on particular trial types (e.g. correct vs. incorrect, or laser on vs off), binning by sensory evidence level was based on all trials.

#### GLM-HMM

To identify sustained stretches of high-performing trials during the accumulation-of-evidence task, we fit behavioral data to a hidden Markov model (HMM) with generalized linear model (GLM) observations governing choice behavior in each state (GLM-HMM; **Extended Data Fig. 3**). A full description of the model as applied to binary decision-making behavior was previously reported^9,44^, and the underlying code can be found at https://github.com/irisstone/glmhmm.

Briefly, the hybrid model is defined by a transition matrix containing a fixed set of probabilities that govern the transitions from one state to another on a given trial, as well as a vector of state-specific GLM weights that map external task covariates onto the probability of a choice (**Extended Data Fig. 3a**). We used the following trial-by-trial task covariates to predict the binary choice (left, −1; right, 1) of mice: a bias term, the normalized difference in the number of right minus left cues (Δ cues, #right-#left), prior trial choice (left, −1; right, 1), prior trial reward (left rewarded, −1; right rewarded, 1; unrewarded, 0), and the presence of laser (off, 0; on, 1).

To identify the number of GLM-HMM states that best explains the behavioral data, we computed the cross-validated log-likelihoods of model fits to data, aggregated across sessions and mice, with indirect or direct pathway inhibition data fit separately (**Extended Data Fig. 3b**). Held out test sets (five folds) consisted of ∼20% of randomly selected sessions with the constraint of having approximately equal numbers of sessions from individual mice, in order to decrease the influence of individual differences to model fits. The remaining ∼80% of sessions were used to fit the model under parameterizations of 1-5 states. Each model was fit 20 times using different initializations of the weights and transition matrix. We initialized state transitions using a Dirichlet distribution (assuming a bias towards within-state transitions trial-by-trial), and we initialized GLM weights based on fits to the 1-state model (assuming each state would have weights in this range). We confirmed that model weights from the top four fits from each initialization did not vary greater than +/-0.05, and selected the best fit for further characterization.

We then expressed the log-likelihood for each model in units of bits per session by subtracting the log-likelihood of the 1-state model and dividing by the number of held out sessions scaled by the binary logarithm,

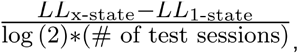

where *LL*_*x-state*_ is the log-likelihood of the model with *x* states.

We utilized a 3-state GLM-HMM based on the plateau in log-likelihoods around ∼3-5 states (**Extended Data Fig. 3b**). State 1 (“engaged”) was characterized by relatively higher weights on sensory evidence (Δ cues) and the laser, and smaller weights on previous choice. In contrast, state 2 (“right-based”) and state 3 (“left-biased”) were characterized by large and opposite weights on the bias term and smaller weights on sensory evidence and the laser (**Extended Data Fig. 3c**). These states were highly persistent, as reflected by the high within-state transition probabilities learned by the model (**Extended Data Fig. 3d**). Choice accuracy was highest in the engaged state (**Extended Data Fig. 3e**), consistent with learned GLM weights. All mice occupied the engaged state as well as at least one of the left or right biased states (**Extended Data Fig. 3f**). Consistent with prior work^9^, the influence of laser was greatest during the engaged state compared to both biased, or “disengaged”, states (**Extended Data Fig. 3c,g**).

### Data Selection Criteria

For all behavioral and neural analysis, we only considered trials in which mice occupied a task-engaged state (see **Behavioral Analyses**, *GLM-HMM*, state 1, “engaged”; ∼62% of all trials). Each trial was assigned to the state with the maximum posterior probability given the inputs and choice data.

From engaged state trials, we also removed trials in which mice traversed the same maze position multiple times by making “early turn” motor errors (i.e. an absolute view angle greater than 90° prior to 290-cm in maze position; ∼9% of “engaged” state trials), as well as “warm-up” maze trials in which a visual guide in the rewarded arm was visible throughout the trial (see **Virtual Reality Behavior**, *Accumulation-of-evidence task*, Maze 4; ∼7% of all trials).

After applying the above trial selection criteria, to ensure individual sessions were sufficiently powered in trials for neural analyses, we only considered sessions with at least 50 laser off trials, and 10 laser on trials. To ensure sufficient power in neurons for population-based analyses (**Fig. 3g-i**; **Fig. 4d-f**; **Extended Data Fig. 6**; **Extended Data Fig. 9c**), we only considered sessions that met the above criteria and also contained greater than 50 simultaneously recorded neurons (with mean firing rates >1 Hz).

After the above data selection criteria, between one and seven recording sessions were included for analysis per mouse. Sessions from the same recording bank for a single mouse did not exceed three (see **Neuropixels Data Acquisition**, *Recording site selection* and *Removal of duplicate clusters*).

### Neural Analyses

#### Preprocessing of on-task recordings

To obtain firing rates in the accumulation-of-evidence task, each neuron’s spikes were first counted in 10 ms bins across the session and then smoothed using a causal half-Gaussian filter with 400 ms standard deviation.

All task-related neural activity was analyzed as a function of position in the central stem of the maze. This involved accounting for the duration mice occupied a given maze position on each trial^7,8^. Specifically, we used the duration in each trial mice occupied y-positions −30-to-300 cm, in binned increments of 5 cm (66 total bins; start region: 6 bins, −30-0 cm; cue region: 40 bins, 0-200 cm; delay region: 20 bins, 200-300 cm). Firing rate (Hz) in each position bin was calculated by first averaging the neuron’s smoothed spike train for the time points spent in each position bin, and then dividing by the duration of time spent in the position bin.

To facilitate comparisons across neurons with baseline firing rates and deviations in firing rate that could vary widely, we z-scored the firing rate of each neuron at each position bin by subtracting the neuron’s mean firing rate across position bins and trials and dividing by the neuron’s standard deviation in firing rate across position bins and trials (**Fig. 1; Fig. 3g-i; Fig. 4d-f; Extended Data Fig. 2c-g; Extended Data Fig. 6; Extended Data Fig. 9c**).

#### Preprocessing of off-task recordings

Since mice did not traverse a virtual maze with spatial positions in off-task recordings, neural activity was processed in temporal, not spatial, bins (ACC activity: **Fig. 1g-h; Extended Data Fig. 2c**; dorsal striatal activity: **Extended Data Fig. 2d,f-g**). Specifically, smoothed spike trains (as performed in *Preprocessing of on-task recordings)* were instead aligned to the 3 s preceding laser delivery up until 3 s following laser offset. This alignment across laser sweeps achieved equally sized smoothed spike trains during baseline (−3-0 s), laser on (0-3 s), and post-inhibition (3-6 s) temporal epochs (**Extended Data Fig. 2a**).

Off-task smoothed spike trains were z-score normalized by subtracting the mean of off-task firing rates prior to laser delivery (baseline, −3-0 s), and dividing by the standard deviation in firing rates on-task (**Fig. 1g-h**; **Extended Data Fig. 2c-g**). The standard deviation in on-task firing rates was used as a more accurate estimate of the variance in firing rate for each neuron given the longer recording duration (>1 hour on-task versus ∼5 minutes off-task).

#### Basic characterization of laser modulation

We used basic descriptive statistics on the averaged normalized firing rates of neurons to assess their modulation by pathway inhibition during on- and off-task contexts (**Fig. 1g-h**; **Extended Data Fig. 2c-g**). In the on-task context, we averaged activity in the cue region – where inhibition was delivered (see **Optogenetics**; **Fig. 1d**) – on all trials. For each neuron, we then used an unpaired two-tailed Mann-Whitney U test comparing cue region activity on laser on versus off trials to assess significant modulation by laser. In the off-task condition, for each neuron we averaged activity in the baseline (−3-0 s) and laser delivery (0-3 s) periods for each laser sweep (30-40). We then used a two-tailed paired Wilcoxon signed rank test to compare activity with and without laser. Neurons with p<0.05 in each task setting and with each statistical test were deemed significantly modulated by laser. The direction of modulation (‘excited’ or ‘inhibited’) was determined by comparing mean activity across all laser on and off trials in each setting.

To assess the significance of the proportions of excited versus inhibited neurons across groups (indirect pathway, direct pathway, or no opsin), and/or across experimental conditions (on-versus off-task), we used two-tailed two-proportion Z-tests on the fraction of significant neurons in each category or group (**Fig. 1h; Extended Data Fig. 2f**).

#### Linear encoding model

To characterize each neuron’s encoding of behavioral variables in the accumulation-of-evidence task, we used a linear encoding model to predict each neuron’s trial-by-trial position-binned firing rate (Hz) based on task variables (**Extended Data Fig. 4**; *generalized linear regression model, fitglm* function in MATLAB). We used the following task variables: (i) previous trial outcome (incorrect, −1; correct, +1), (ii) sensory evidence (#ipsilateral - #contralateral cues up to each position bin; contralateral, −1; ipsilateral, 1), (iii) current trial choice (contralateral, −1; ipsilateral, +1), and (4) the presence of laser (off, −1; on, +1). From each neuron and each position bin, we obtained estimated coefficients reflecting the relative weighting of these variables in predicting neural activity at each position bin across trials.

For initial characterization of all ACC (**Fig. 2**) and dorsal striatal (**Extended Data Fig. 5**) neurons, the laser predictor was omitted from model fits and only laser off trials were used for encoding models. For characterizing the effect of pathway-specific inhibition based on each neuron’s tuning profile (**Fig. 3d-f; Fig. 4a-c; Fig.5a-c; Extended Data Fig. 7; Extended Data Fig. 9a-b**), the laser predictor was included in model fits and both laser off and on trials were considered.

#### Statistical definition of task-relevant neurons

From each encoding model fit, we obtained a t-statistic for each task variable at each position bin, which reflects the significance of this variable’s influence based on the null hypothesis that the coefficient is zero. To assess if a neuron was significantly tuned to a given task variable across any spatial bin in the cue or delay region, we had to appropriately control for multiple hypothesis testings. To this end, for each neuron and task variable, we compared the maximum and minimum t-statistics across position bins to a null distribution of t-statistics obtained from shuffled data. Shuffled data preserved the temporal order of trial-by-trial binned firing rates, but randomly reordered the trial-by-trial prediction matrix of task variables (repeated 200 times). For each neuron and task variable, we then obtained two null distributions consisting of either the maximum or minimum t-statistics for all 200 fits to randomly shuffled data. A neuron was then termed significantly tuned to a task variable if the maximum t-statistic from the real data for a given variable exceeded the 95th percentile of that in the null distribution of maximum values, or if the minimum t-statistic from the real data was smaller than the 5th percentile of that in the null distribution of minimum values.

Some neurons with significant evidence encoding also exhibited significant choice encoding (**Fig. 2d**; **Extended Data Fig. 4**). We categorized a neuron as ‘evidence-tuned’ if it showed significance for the evidence coefficient at any maze position (**Fig. 2-5; Extended Data Fig. 4-5,9**), and ‘choice-tuned’ if it showed significance for the choice, but not evidence, coefficient at any position (**Fig. 2 and 5; Extended Data Fig. 4-5,9**).

The ipsilateral and contralateral preference of each neuron was determined by the averaged evidence or choice coefficient across cue and delay maze positions, with a positive value indicating ipsilateral preference and a negative value indicating contralateral preference.

#### Model-based characterization of laser effects

We utilized the encoding model’s coefficients to analyze the influence of pathway inhibition on neural activity using the following approaches.

First, we examined correlations between evidence and laser coefficients (**Fig. 3d**; **Fig. 4a**), or choice and laser coefficients (**Fig. 5a**), to assess patterns between the direction of laser influence and the side preference of evidence or choice-tuned neurons. Toward this end, for each neuron we averaged the evidence (or choice) coefficient and the laser coefficient across the cue and delay regions. We used Spearman rank correlation (*corr* function in MATLAB, with ‘Spearman’ option) to then obtain cross-neuron correlation values and assess their significance.

Second, we visualized laser coefficients averaged across neurons at each position in the maze (**Fig. 3e**; **Fig. 4b**), separately for neurons grouped by the sign and significance of their evidence coefficients (see *Statistical definition of task-relevant neurons)*. For comparison, we visualized the laser coefficients at each position averaged across neurons that did not show significance for evidence or choice coefficients (“un-tuned”, or “non-decision-tuned”).

Third, we isolated neurons based on their tuning profile (evidence-, choice-, or non-decision-tuned) and side preference (ipsilateral or contralateral) using the statistical definition described above (see *Statistical definition of task-relevant neurons*), and constructed population-averaged evidence tuning curves based on their firing rates, separately for laser off and on trials (see *Evidence tuning curves for single neurons* below for further detail; for ACC, **Fig. 3f**; **Fig. 4c**; **Fig. 5b**; for striatum, **Extended Data Fig. 9a-b**).

Finally, we directly compared the laser coefficients averaged across the cue and delay regions for neurons grouped by the sign and significance of their evidence and choice coefficients (**Fig. 5c**). To assess statistical significance, we used a 2-way analysis of variance (ANOVA) of averaged laser coefficients with tuning profile (evidence-, choice-, or non-tuned; separately for ipsilateral- and contralateral-preferring neurons) and pathway inhibition (direct or indirect) as factors. Where significant interactions were observed (tuning x pathway, p<0.05), we used *post hoc* unpaired t-tests to compare laser coefficients between each pair in the group.

#### Evidence tuning curves for single neurons

To visualize the dependence of neural activity on varying levels of evidence (**Fig. 2b, 3f, 4c and 5b; Extended Data Fig. 4d-f, 5a, 7, and 9a-b**), we constructed evidence tuning curves across neuron classes identified with our encoding model. First, subpopulations of neurons were pooled together according to the significance and sign of their coefficients (see *Statistical definition of task-relevant neurons*), and trials were divided based on the final evidence value (see **Behavioral Analyses**, *Psychometric behavior and binning of sensory evidence levels*).

To enable cross-neuron comparison, firing rates of each neuron were normalized by subtracting the mean firing rate at each position bin, and dividing by the standard deviation across all position bins and trials. Normalized activity was then averaged across trials in each bin of sensory evidence. Averages were made at each position bin in order to visualize evidence-tuned activity along the maze (**Fig. 2c; Extended Data Fig. 4d-h and 5b**) or across all positions in the cue and delay regions to construct evidence tuning curves (**Fig. 2b, 3f, 4c and 5b; Extended Data Fig. 4d-f, 5a, 7, and 9a-b**).

Given the ipsilateral or contralateral preference of evidence-tuned neurons were consistent across correct and incorrect trials (**Fig. 2a-c**; *top*), both trial types were used for constructing tuning curves for evidence-tuned neurons (**Fig. 3f and 4c; Extended Data Fig. 9a**). As choice-tuned neurons showed inverted ipsilateral and contralateral preferences on correct versus incorrect trials (**Fig. 2a-c**; *bottom*), tuning curves were constructed separately for correct and incorrect trials (**Fig. 5b; Extended Data Fig. 9b**).

#### Statistical assessment of laser effects in task-relevant neurons

To assess significance of laser modulation across the population of evidence-tuned neurons (**Fig. 3f and 4c; Extended Data Fig. 9a**), we fit a linear mixed-effects model (*fitlme* function in MATLAB) to predict the normalized neural activity averaged across the cue and delay regions based on the final evidence value of a trial (normalized to [0,1] per session) and laser delivery (0 for off, 1 for on trials) as fixed effects, and random effects for intercept, evidence and laser grouped by individual neurons (based on both correct and incorrect trials): *neural activity ∼ 1 + evidence + laser + (1 + evidence + laser* | *individual neuron)*. Specifically, we used the p-value associated with the laser term (using the t-statistics for testing the null hypothesis that the coefficient is equal to zero), to determine if the activity of evidence-tuned neurons with a given side preference were significantly upward-(indicating excitation; positive laser coefficient) or downward-shifted (indicating inhibition; negative laser coefficient) with pathway-specific inhibition. The same processing and statistical test were used for neurons that were tuned to neither evidence nor choice (**Extended Data Fig. 7**).

To test if the population of choice-tuned neurons show significant modulation by laser (**Fig. 5; Extended Data Fig. 9b**), we fit a similar linear mixed-effects model, but with choice (0 for contralateral, 1 for ipsilateral choice trials) instead of evidence (based on both correct and incorrect trials): *neural activity ∼ 1 + choice + laser + (1 + choice + laser* | *individual neuron)*.

We focused on the p-value associated with the laser term (using the t-statistics for testing the null hypothesis that the coefficient is equal to zero), to determine if the activity of choice-tuned neurons with a given side preference were significantly affected by the pathway-specific inhibition.

#### Obtaining the evidence decoder axes

To find the direction of the neural population activity in each recording session that best decodes evidence (‘decision axis’) (**Fig. 3g-i; Fig. 4d-f; Extended Data Fig. 6 and 9c**), we applied linear regression with lasso (L1) regularization to the activity of all simultaneously recorded neurons in each session (see **Data Selection Criteria** for session inclusion criteria). We obtained a separate decoder for each position. The response variable was the cumulative evidence (#ipsilateral - #contralateral cues) up to each position across trials, which was normalized to [-1,1]. Predictor variables were the z-scored neural activity (see *Preprocessing of on-task recordings)* of all simultaneously recorded neurons in a session. Only trials without laser delivery were used for training (both correct and incorrect trials). We used 5-fold cross-validation to assess the predictive performance of decoders and to obtain neural projections (in other words, decoder axes were projected to untrained portions of neural data; see *Projection of population activity to the evidence decoder axis* below). Since we used 5-fold cross-validation, this produced a set of 5 decoder axes (five *n* neurons by 1 unit vectors) at each position. In order to find optimal regularization parameters λ, within each of these 5-folds, we used another inner-layer 5-fold cross-validation (“nested cross-validation”) from each training fold^94,95^.

For each session, we performed the following steps:

1. All laser off trials were first split into 5 folds (‘outer split’). To optimize the generality of model fits to each fold, trials were pseudo-randomly distributed such that each fold sampled a similar distribution of evidence values. Each split (20% of laser off trials) was assigned as ‘test’ trials and the remaining ones (80% of laser off trials) were assigned as ‘training’ trials.
2. To find an optimal regularization parameter with nested cross-validation, each of the 5 training folds was further split into 5 folds (‘inner split’) and similarly assigned to inner test and training trials. For each inner training fold, we trained regression models with 15 different λ parameters (logarithmically scaled from 0 to 1) and evaluated their model performance with each corresponding inner test block, using Pearson’s correlation coefficients between actual and predicted evidence as a performance metric^8,43^.
3. Step (ii) was repeated for all inner splits, and the λ value that yielded the best performance across 5 sub-fold iterations was selected for regularization.
4. We trained a single regression model with the selected λ value, and evidence values and neural data of an outer training fold, to obtain a single decoder axis **w**_**1**_ (*n* neurons x 1 vector) at each position. Each decoder axis was normalized by dividing by the Euclidean norm of vector weights^96^.
5. Performance of this single regression model with ‘optimal’ λ was evaluated on the held-out test trial fold (**Extended Data Fig. 6b**, ‘data’), using Pearson’s correlation coefficients between actual and predicted evidence.
6. We repeated steps (ii) to (v) for the other 4 outer folds to complete a set of five n x 1 decoder axes, **w**_**1-5**_ for each position.
7. Steps (i) to (vi) steps were then repeated to acquire a set of evidence decoder axes across 60 positions that span the cue and delay regions.

#### Assessing decoder model performance

To show that regression models can decode the level of accumulated evidence from population neural activity above chance levels, we compared the decoding accuracy of regression models from real data to those obtained from shuffled data. Shuffled data preserved the temporal order of projected activity across position bins, as well as the progression of cumulative sensory evidence across position within a trial, but randomly permuted their relationship. Across ten randomly shuffled data sets, we repeated steps (i) to (vii), calculated the average decoding accuracy across each fit to shuffled data, and compared this to the decoding accuracy when fit to real data (**Extended Data Fig. 6b**, ‘data’ vs. ‘shuffled’).

#### Projection of population activity to the evidence decoder axes

After we obtained decoder axes **w**_**1-5**_ at each position, we projected the z-scored neural data of the outer fold test trials **X**_**1**_ (N_1_ trials x *n* neurons) at each position onto the corresponding decoder axis **w**_**1**_ (*n* neurons x 1) by taking the dot product, to get the cross-validated population projection **p**_**1**_ (N_1_ trials x 1). This step was repeated for neural data of other 4 outer fold test trials **X**_**2-5**_ to have **p**_**1-5**_ (N_1-5_ trials x 1). For laser trials, which were not used for acquiring decoder axes, we projected the neural activity at each position to the corresponding decoder axis from 5 folds at each position, and then averaged the results across the 5 iterations. These projection steps were repeated across 60 positions.

#### Visualization of evidence projections and their laser modulation

When visualizing time-varying projections onto the evidence axes on laser off trials (**Fig. 3g and 4d; Extended Data Fig. 6c and h**), the evidence-axis projected neural data were first smoothed for each trial using a causal half-Gaussian filter with standard deviation of 10 cm. Then the mean of laser off trials at each position was subtracted from the projected activity to facilitate cross-session comparison. Evidence-axis projected activity was then separated into 6 trial bins based on their final evidence values (see **Behavioral Analyses**, *Psychometric behavior and binning of sensory evidence levels*), selecting only laser off trials from each bin. To visualize neural trajectories averaged across trials, we grouped trials based on the final evidence level, rather than current or cumulative evidence at each maze position, in order to avoid individual trials being assigned to different bins of sensory evidence at different positions. This facilitated the visualization of trial-level neural trajectories.

To assess if pathway-specific inhibition causes overall shifts in neural trajectories toward greater ipsilateral or contralateral coding (**Fig. 3h and 4e**), we calculated the mean neural trajectories on laser off versus on trials from each session, and visualized the average and s.e.m. across sessions.

#### Evidence tuning curves for evidence-axis projections

To examine the dependence of evidence-axis projections on varying levels of evidence and to visualize the impact of pathway inhibition on them, we constructed evidence tuning curves for evidence axis-projected neural activity (**Fig. 3i and 4f**, *left*, for ACC**; Extended Data Fig. 9c**, *left*, for dorsal striatum). First, the mean of laser off trials were subtracted from unsmoothed projections at each maze position. We then averaged projected activity across the cue region in trials binned by their final evidence levels (see **Behavioral Analyses**, *Psychometric behavior and binning of sensory evidence levels*), separately for laser on and off trials. We then took the average and s.e.m. of these values across sessions.

We focused on the cue region for this analysis. In the cue region, on both correct and incorrect trials, the evidence-axis projections showed a similar overall pattern (**Extended Data Fig. 6c,h**, *right*, black), consistent with the tuning properties of evidence-tuned neurons on correct and incorrect trials at the single neuron level (**Fig. 2b**, *top*, for ACC; **Extended Data Fig. 5a**, *top*, for striatum). This preserved pattern across correct and incorrect trials validated using both trial types in this analysis. Doing so was helpful for interpreting the results, given that the laser can generate incorrect trials, and therefore omission of incorrect trials would amount to omitting trials where the laser had an effect. In contrast, in the delay region, evidence-axis projections reversed their side preferences on incorrect trials (**Extended Data Fig. 6c,h**, *right*, gray), an effect likely driven by the choice-tuned neurons (**Fig. 2c**). This is consistent with choice-selectivity being most prominent towards the end of the central maze stem (**Fig. 2c**).

#### Statistical assessment of laser modulation in evidence projections

To statistically test for shifts in evidence-axis projections by pathway inhibition (**Fig. 3i and 4f**, *left*, for ACC**; Extended Data Fig. 9c**, *left*, for dorsal striatum), we fit a linear mixed-effects model to predict the cue period averaged evidence-axis projected activity across sessions (*fitlme* function in MATLAB). For each pathway inhibition group, we used the final evidence value on each trial (normalized to [0,1]) and laser delivery (0, off; 1 on) as fixed effects, and random effects for intercept, evidence, and laser for each session: *evidence projection ∼ 1 + evidence + laser + (1 + evidence + laser* | *individual session)*. We reported the p-value associated with the laser term (using the t-statistics for testing the null hypothesis that the coefficient is equal to zero), to determine if the evidence projections were significantly affected by pathway-specific inhibition.

#### Relationship between the neural and behavioral shifts induced by laser

To probe the correlation between the shift in evidence-axis projection population activity and the decision bias induced by pathway-specific inhibition at the level of sessions (**Extended Data Fig. 6g and l**), we first averaged the unsmoothed neural projections, separately for laser off and on trials, across the second half of the cue region from each session. The neural shift induced by pathway inhibition was defined as the difference between these two mean values (on - off). We used laser-induced ‘choice bias’ (see **Behavioral Analyses**, *General performance*) as an index for behavioral shift by the laser. These two variables were tested with Pearson’s correlation separately for inhibition of each pathway.

#### Obtaining the first principal component axis

To examine if pathway-specific inhibition shifted activity along the axis that explains the most variance in population activity, we first obtained the first principal component (PC1) axis from trial-averaged population activity. This dimension of population activity primarily reflects the progression of time or position within a trial (**Extended Data Fig. 6d-f,i-k**). We took a similar approach as explained above (see *Obtaining the evidence decoder axis)* for separating data into test and training sets. For trials from each of the outer training folds, we prepared a neural data matrix, where each row is the trial-averaged, z-scored firing rate of a neuron across the cue region (*n* neurons x 40 position bins). We centered each column of this matrix and performed the principal component analysis (*pca* function in MATLAB) along the neuron dimension, to find a PC1 axis for each session with weights on the *n* neurons. The PC1 axis was normalized by dividing by the Euclidean norm of the vector. These steps were repeated for the remaining outer training folds to acquire a set of five PC1 axes per session.

#### Projection of population activity to the first principal component axis

After obtaining PC1 axes, the population neural activity of held-out laser off trials were projected to the corresponding PC1 axis, repeatedly at each position. Population activity of the laser on trials at each position was projected to each of the PC1 axes and then averaged across 5 iterations. The same projection for laser on trials was repeated across positions.

PC1 projections across outer training folds both within and across sessions showed highly similar and stereotyped profiles – typically, a brief activity increase (in arbitrary units) and then progressive decrease during the cue region (**Extended Data Fig. 6d,i**). However, the polarity of this profile was occasionally reversed. To ensure the consistency of PC1 projections across outer training folds within a session, we calculated the Pearson’s correlation coefficient between the PC1 axis from the first fold to that of all other folds, and multiplied the weights by −1 if the correlation value was negative. To ensure the consistency of the sign of PC1 projections across sessions, we multiplied by −1 if the trial-averaged PC1 projections increased between the beginning versus end of the cue region.

#### Evidence tuning curves for the first principal component axis projections

Evidence tuning curves (**Fig. 3i and 4f**, *right*) were similarly constructed for PC1-axis projections as for evidence-axis projections (see *Evidence tuning curves for evidence projections)*. We averaged projected activity across the cue region in trials binned by their final evidence levels (see **Behavioral Analyses**, *Psychometric behavior and binning of sensory evidence levels*), separately for laser on and off trials. We then took the average and s.e.m. of these values across sessions.

#### Statistical assessment of laser modulation in the first principal component axis projections

The same linear mixed-effects model described in *Statistical assessment of laser modulation in evidence projections* was used to predict laser-induced shifts in PC1-axis projected activity (**Fig. 3i and 4f**, *right*, for ACC; **Extended Data Fig. 9c**, *right*, for dorsal striatum) in the cue region: *PC1 projection ∼ 1 + evidence + laser + (1 + evidence + laser* | *individual session)*. We reported the p-value associated with the laser term (using the t-statistics for testing the null hypothesis that the coefficient is equal to zero), to determine if the evidence projections were significantly shifted by the pathway-specific inhibition.

#### Decoding the maze position from the first principal component axis projections

Population projections to the PC1 axis appeared to correlate with the progression through position in the cue region (**Extended Data Fig. 6d,i**). To further confirm that PC1-projected population activity indeed contained information about position in the maze, we used a linear regression model (5-fold cross-validation, laser off trials only) to decode mouse position in the maze on each trial based on PC1-projected population activity. Model performance was assessed by calculating the Pearson’s correlation coefficient between actual and predicted positions. To confirm if decoding position from PC1 projections was better than chance, we repeated the regression from data where position was randomly shuffled, evaluated the mean performance across 10 different shuffled data sets, and compared with the decoding performance between shuffled and unshuffled data using a two-tailed Mann-Whitney U test.

### Statistics and Reproducibility

No statistical methods were used to predetermine sample sizes, but they were chosen based on previous studies that used comparable techniques or experiments^4,54,62,97^, and on the availability of transgenic animals. Mice were allocated to groups according to their strains. Experimenters were not blinded to mouse group allocation during experiments and analyses. All results were successfully replicated across multiple sessions and cohorts of mice. We used multiple independent analyses within each experiment, to confirm our findings whenever possible. Common analysis pipelines were used in all cases, without the need for manual intervention. All statistical tests were performed with MATLAB (2023b; Mathworks) or Prism (version 9.5.1; GraphPad). Details on specific tests performed are included in appropriate sections in **Behavioral analyses** and **Neural analyses**. All statistical test outputs – such as sample sizes, exact p-values, test-statistics, and degrees of freedom – will be included in Supplementary Table 1.

### Model Simulations

#### Simulated hypotheses of interactions between evidence- and choice-tuned neurons

In **Extended Data Figure 8**, we simulated three hypotheses for interactions between indirect pathway inhibition, ACC evidence-tuned neurons and choice-tuned neurons, and action for the effects of the pathway manipulation. For each hypothesis, we used a subset of approximately 9000 trials from experimental indirect pathway sessions to generate the number of left and right cues, with the same statistics as the engaged trials we analyze in the neural data. Due to the probabilistic outcomes of our simulations, we simulated each of these real trials four times for each hypothesis to generate the simulated dataset.

*Hypothesis 1i: Choice neurons as immediate readout of evidence neurons, with incorrect trials driven by noise in the choice neurons*

Data for hypothesis 1i (**Extended Data Fig. 8a-d**) was generated according to the following model. For a trial with final evidence *e* (the difference between the number of right and number of left cues), the firing rates, *e*_*i*_ and *e*_*c*_, of the ipsilateral- and contralateral-preferring evidence-tuned neurons respectively were given by:

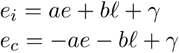

where a = 1, b = −2, *γ* = 10, and *ℓ* is an indicator variable that takes value 1 if the laser was on in a trial.

We simulated a population of 10 ipsilateral-preferring and 10 contralateral-preferring choice-tuned neurons. The firing rates, *c*_*i*_ and *c*_*c*_, of each of the ipsilateral- and contralateral-preferring choice neurons, respectively, were independently simulated according to:

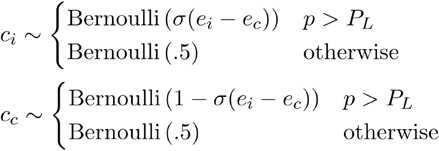

where *p* is random uniform variable, drawn independently for each choice-tuned neuron for each trial, *P*_*L*_ = 0.5 is the probability of an individual choice neuron lapsing, and *σ* is a logistic function parameterized as

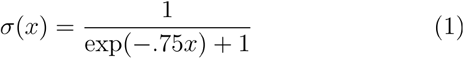

The action *A* was determined by:

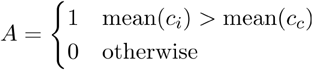

where *A* = 1 corresponds to an ipsilateral choice and *A* = 0 corresponds to a contralateral choice.

*Hypothesis 1ii: Choice neurons as immediate readout of evidence neurons, with incorrect trials driven by noise in the evidence neurons*

Data for hypothesis 1ii (**Extended Data Fig. 8a,e-g**) was generated according to the following model. To simulate noise in the evidence-tuned neurons, we assumed that an animal fails to attend to a fraction *d* =.3 of the cues since previous analyses of this task have suggested that sensory noise is the dominant noise source in the behavior^98^. For each cue in the set of left cues *L*, we drew *z*, a random uniform number between 0 and 1, and if *z* > *d*, we added the cue to the attended set of left cues *L’*. We repeated the same process for each cue in the set of right cues *R* to generate *R’*. We define *e’* as the difference between the number of cues in *R’* and *L’*. The firing rates, *e*_*i*_ and *e*_*c*_, of the ipsilateral- and contralateral-preferring evidence neurons, respectively, were given by:

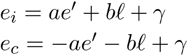

where a = 1, b = 2, *γ* = 10, and *ℓ* is an indicator variable that takes value 1 if the laser was on in a trial.

The firing rates, *c*_*i*_ and *c*_*c*_, of each of the ipsilateral- and contralateral-preferring choice neurons respectively were independently simulated according to:

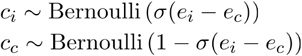

where *σ* is the logistic function in Eq. (1).

The action *A* was determined identically to Hypothesis 1i.

*Hypothesis 2: Choice neurons as action readout*

Data for hypothesis 2 (**Extended Data Fig. 8h-k**) was generated according to the following model. For a trial with final evidence *e* (the difference between the number of right and number of left cues), the firing rates, *e*_*i*_ and *e*_*c*_, of the ipsilateral- and contralateral-preferring evidence-tuned neurons respectively were given by:

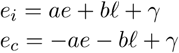

where a = 1, b = −2, *γ* = 10, and *ℓ* is an indicator variable that takes value 1 if the laser was on in a trial.

The action *A* was probabilistically drawn according to:

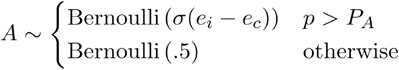

where *p* is a uniform random variable drawn independently for each trial, *P*_*A*_ = 0.1 is the probability of a behavioral lapse, and *σ* is the logistic function in Eq. (1).

The firing rates, *c*_*i*_ and *c*_*c*_, of each of the ipsilateral- and contralateral-preferring choice-tuned neurons, respectively, were independently simulated according to:

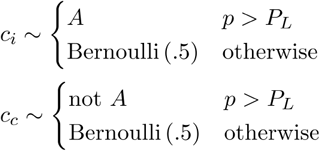

where *p* is a uniform random variable drawn independently for each neuron on each trial and *P*_*L*_ = 0.1 is the lapse rate of the choice-tuned neurons reading from the action.

*Hypothesis 3: Choice neurons as multi-process readout of either evidence neurons or an alternative decision process*

Data for hypothesis 3 (**Extended Data Fig. 8l-o**) was generated according to the following model. For a trial with final evidence *e* (the difference between the number of right and number of left cues), the firing rates, *e*_*i*_ and *e*_*c*_, of the ipsilateral- and contralateral-preferring evidence-tuned neurons respectively were given by:

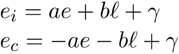

where a = 1, b = −2, *γ* = 10, and *ℓ* is an indicator variable that takes value 1 if the laser was on in a trial.

The firing rates, *c*_*i*_ and *c*_*c*_, of each of the ipsilateral- and contralateral-preferring choice-tuned neurons respectively were given by:

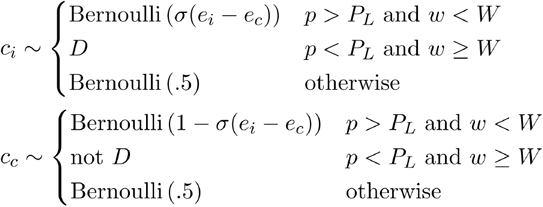

where *w* is random uniform variable, drawn independently for each trial but is the same for all choice neurons, *W* = 0.5 is the probability that the choice neurons readout evidence rather than the alternative decision process on a given trial, *D* ∼ Bernoulli(.5) is the value of the alternate decision process drawn independently on each trial but takes the same value for all neurons reading out from the alternative decision process, *p* is a uniform random variable drawn independently for each choice neuron for each trial, *P*_*L*_ = 0.5 is the probability of an individual choice neuron lapsing, and *σ* is the logistic function defined in Eq. (1). The term “not D” is defined as 1 if D = 0 and 0 if D = 1, such that the response of the contralateral cells is consistent with the ipsilateral cells to the alternative decision process.

The action A was determined by:

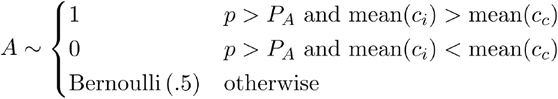

where *A* = 1 corresponds to an ipsilateral choice and *A* = 0 corresponds to a contralateral choice, *p* is a uniform random variable drawn independently on each trial, and *P*_*A*_ = 0.1 is the probability that the behavior lapses from the choice neuron readout.

### Histology

All mice were anesthetized with a 0.05 mL injection of Euthasol (intraperitoneal) and transcardially perfused with phosphate-buffered saline (PBS) followed by 4% paraformaldehyde (PFA) for tissue fixation. Decapitated heads with intact fiberoptic implants were post-fixed in 4% PFA for ∼2 h, followed by brain extraction and 24 h of additional post-fixation in PFA. Brain samples were then stored in PBS. Described in greater detail below, brains were then rendered transparent using a previously described technique for immunolabeling-enabled three-dimensional imaging of solvent-cleared organs (iDISCO+)^99,100^, imaged on a fluorescent light-sheet microscope (Life Canvas Technologies, SmartSPIM) to identify CM-DiI-labeled fiberoptic and electrode tracks as well as viral expression of GFP, and then registered to the Allen Common Coordinate Framework.

#### Tissue preparation

Briefly, similar to previous iDISCO+ procedures^99–101,92^, brains were first serially dehydrated in increasing concentrations of methanol (Carolina Biological Supply, 874195; 20, 40, 60, 80, 100% in doubly distilled water (ddH_2_0); 45 min - 1 hr duration each) followed by a final incubation in 100% methanol overnight. Brains were then bleached in 5% hydrogen peroxide (Sigma H1009) in methanol overnight, followed by serial rehydration in decreasing concentrations of methanol (100, 80, 60, 40, 20% in ddH_2_0; 45 min – 1 hr duration each). Finally, brains were washed in Dulbucco’s PBS (DPBS, Thermo Fisher Scientific; 1x 45 min - 1 hr), permeabilized with 0.2% Triton X-100 (Sigma T8787) in PBS (PBS-Tx; 2x 45 min - 1 hr), and then incubated in 20% DMSO (Fisher Scientific D128) + 0.3 M glycine (Sigma 410225) PBS-Tx (1x 45 min - 1 hr). All dehydration, rehydration and washing steps were performed at room temperature.

#### Whole-brain immunolabeling and clearing

Endogenous epitopes were first blocked by washing brains in 10% DMSO + 6% normal donkey serum (NDS; EMD Millipore S30) in PBS-Tx for 2-3 days. Next, brains were washed in PTwH solution (PBS + 0.2% Tween-20 (Sigma, P9416) + 10-µg/mL heparin; 2x for 1 hr) before incubating with primary antibody (chicken anti-GFP, 1:500; Aves GFP-1020) solution (5% DMSO + 3% NDS in PTwH) for 7 days. Following primary antibody incubation, brains were washed in PTwH solution six times at increasing durations (10, 15, 30, 60, 120 min, and then overnight), and then incubated in secondary antibody (Alexa Fluor 647 donkey anti-chicken, 1:500; Jackson ImmunoResearch; code: 703-6-6-155) solution (3% NDS in PTwH) for 7-days. Following secondary antibody incubation, brains were once again washed in PTwH six times at increasing durations (10, 15, 30, 60, 120 min, and then overnight). Finally, brains were serially dehydrated in increasing concentrations of methanol (20%, 40%, 60%, 80%, 100% in ddH20; 45 min – 1 hr duration each), and then cleared for imaging by incubating in a 2:1 solution of dichloromethane (DCM; Sigma 270997):methanol for 3 hr followed by two 15 min washes in 100% DCM. Brains were stored long-term in the refractive index-matching solution dibenzyl ether (DBE; Sigma 108014) prior to undergoing light sheet microscope imaging. All blocking, washing, and antibody incubations were performed at 37°C.

#### Light-sheet fluorescence microscopy

Cleared and immunolabeled whole brains were fixed either dorsal- or ventral-side down to a 3D-printed holder with a minimal amount of glue (Loctite, 234796) and immersed in an imaging chamber containing dibenzyl ether. To image cleared whole brain tissue we used a dynamic axial-sweeping light-sheet fluorescence microscope^102^ (Life Canvas Technologies, SmartSPIM) and SmartSPIM acquisition software (https://lifecanvastech.com/products/smartspim/; v5.6). Images were acquired using a 3.6x (0.2 NA) objective with a 3,650 × 3,650 μm field of view onto a 2,048 × 2,048 pixel sCMOS camera (pixel size, 1.78 × 1.78 μm). Horizontal light sheet sections were spaced 2 µM apart. Imaging of the entire brain required 4 × 6 tiling of the horizontal plane across a total of 3,300-3,900 horizontal planes. Images were acquired using 488 nm (autofluorescence channel for brain structure), 561-nm (CM-DiI channel for fiberoptic and electrode shank trajectories), and 647 nm (for anti-GFP labeling of NpHR3.0 viral expression) excitation light at ∼20-90% power (max output: 70 mW) and 2-ms of exposure time.

After acquisition, tiled images from the autofluorescent channel were first stitched into a single imaging volume using the TeraStitcher C++ package^103^ (https://github.com/abria/TeraStitcher). These stitching parameters were then directly applied to the tiled CM-DiI and anti-GFP channel images, which produced three aligned 3D imaging volumes with the same final dimensions. After tile stitching, striping artifacts were removed from each channel using the Python package Pystripe^104^ (https://github.com/chunglabmit/pystripe; v.0.2.0).

#### Atlas registration

To register whole-brain light-sheet microscope images to the Allen Common Coordinate Framework (CCF), we utilized an approach and software developed by the International Brain Laboratory (https://github.com/int-brain-lab/iblapps/wiki)^40^. Briefly, the 488 nm autofluorescence brain volume was first registered to the CCF using the Python package Brainreg (https://github.com/brainglobe/brainreg; v.0.4.0)^105^, and the tissue-to-atlas transformations of this channel were directly applied to the 561 nm CM-DiI and 647 nm anti-GFP volumes. CM-DiI labeled electrode shanks and fiberoptic trajectories were then manually annotated in atlas-registered volumes using the Brainreg-segment Napari module (https://github.com/brainglobe/brainglobe-segmentation; v.0.2.16).

We then used IBL’s Ephys Alignment GUI to refine channel localization in atlas space for each registered Neuropixels 2.0 shank. To do this, we used the atlas registered trajectory of each shank and the ∼10 m baseline recordings we obtained from the most ventral 768 recording sites (spanning 5.76 mm) on each shank (see **Neuropixels Data Acquisition**, *Recording site selection*). The Ephys Alignment GUI provided a default anatomical annotation associated with each registered shank trajectory. We then used the electrophysiological features recorded on each shank at each depth to anchor particular recording sites at precise locations. The principal landmarks we used were reduced single and multi-unit spiking when shank trajectories passed through the corpus callosum, the lateral ventricle, the cell-poor cortical midline, and the surface of the brain. Based on these adjusted anchor points, the remaining electrode locations were determined by spatial interpolation according to the known interelectrode spacing (15 μm). Every recording site was thus localized to a corresponding coordinate and brain area according to the 2017 Allen mouse CCF atlas (https://download.alleninstitute.org/informatics-archive/current-release/mouse_ccf/annotation/ccf_2017/).

#### Spatial organization of task-relevant neurons

To probe the relationship between the anatomical location of recorded neurons and their firing properties, we utilized the *x*-, *y-*, and *z-*axis coordinates (ML, AP, DV) of all recorded neurons and fiberoptic tracks obtained from whole brain registration to the Allen CCF (see above; **Extended Data Fig. 1**). Specifically, to examine a relationship between evidence- or choice-tuning and anatomical position within the ACC or dorsal striatum (**Extended Data Fig. 1c-d**), we plotted the proportion of neurons significantly tuned to each variable as a function of their AP, DV, or ML coordinate separately. For each anatomical axis, we divided the full range of coordinates into four equally-spaced bins to facilitate visualization. To test for statistical significance, we ran logistic regression (MATLAB *fitglm* function with ‘binomial’ distribution) using each neuron’s AP, DV, or ML coordinate to predict if a neuron had significant evidence-tuning or not, or significant choice-tuning or not (tuned, 1; non-tuned, 0). Coordinates were normalized to [0,1] at each spatial dimension. Estimated coefficients and *p*-values for each predictor variable were obtained from the output of the *fitglm* function assuming a two-sided test with the null hypothesis that the coefficient is zero.

### Sex as a Biological Variable

We included both male and female mice (indirect pathway inhibition with A2a::Cre: n = 2 males and 4 females; direct pathway inhibition with D1::Cre: n = 3 males and 3 females; no opsin with A2a::Cre = 2 males). Data from both sexes were pooled together for all analyses.

### Excluded Animals

We excluded one mouse (direct pathway, d1_211) who successfully completed shaping, but failed to perform the accumulation-of-evidence task following electrode implantation surgery. We excluded one mouse (indirect pathway, a2a_245) from all behavioral and neural analyses due to the observation of halorhodopsin expression in both left and right hemispheres.

## Data Availability

All source data for generating figure plots, and processed behavioral and neural data will be made freely available upon publication.

## Code Availability

All code used in the paper will be publicly available on GitHub upon publication.

## Acknowledgements

We thank Mark Goldman, Carlos Brody, and members of the Witten lab and BRAINCoGS U19 team for helpful discussion and critical feedback on the conceptualization of this manuscript; Tim Harris and Wade Sun for the design, manufacture, and attachment of Neuropixels probe dovetail adaptors; Esteban Engel, Oliver Huang, Angela Chan and the PNI Viral Core facility for AAV production; Adrian Sirko and staff at the Princeton Laboratory Animal Resources for help with animal husbandry; Sam Wang and members of his lab for technical assistance in whole brain tissue clearing and light sheet fluorescence microscopy. Funding for this work was provided by NIH U19-NS104648 (I.B.W.), NIH P50-MH136296 (I.B.W.), NIH DP1-MH136573 (I.B.W.), NIH U19-NS123716 (I.B.W.), the Simons Collaboration on the Global Brain (I.B.W.), the Howard Hughes Medical Institute (I.B.W.), CV Starr Fellowship (J.R.C), NIH F32MH118792 (S.S.B.), NIH F32MH132179 (L.S.B.), the Sloan Swartz Foundation (L.S.B.), and the Burroughs Welcome Fund (L.S.B.).

## Author Contributions

J.R.C., S.S.B. and I.B.W. conceived the project and designed the experiments. S.S.B., with assistance from J.R.C., M.S., Y.E., B.M., R.N.F., C.A.Z., A.P.V., M.S., A.G.B., J.L.L., A.L, A.S.K., L.L., and Y.L., performed the experiments. S.S.B., J.R.C., M.S., and T.E. performed analyses on behavioral data. J.R.C. and S.S.B, with inputs from L.S.B., performed analyses on neural data. L.S.B. performed model simulations. I.B.W. obtained funding and supervised the project. J.R.C., S.S.B., L.S.B., and I.B.W. wrote the paper with input from other authors.

## References

1. Alexander, G. E., DeLong, M. R. & Strick, P. L. Parallel Organization of Functionally Segregated Circuits Linking Basal Ganglia and Cortex. Annu. Rev. Neurosci. 9, 357–381 (1986).

2. Albin, R. L., Young, A. B. & Penney, J. B. The functional anatomy of basal ganglia disorders. Trends Neurosci. 12, 366–375 (1989).

3. DeLong, M. R. Primate models of movement disorders of basal ganglia origin. Trends Neurosci. 13, 281–285 (1990).

4. Oldenburg, I. A. & Sabatini, B. L. Antagonistic but Not Symmetric Regulation of Primary Motor Cortex by Basal Ganglia Direct and Indirect Pathways. Neuron 86, 1174–1181 (2015).

5. Lee, H. J. et al. Activation of direct and indirect pathway medium spiny neurons drives distinct brain-wide responses. Neuron 91, 412–424 (2016).

6. Pinto, L. et al. An Accumulation-of-Evidence Task Using Visual Pulses for Mice Navigating in Virtual Reality. Front. Behav. Neurosci. 12, (2018).

7. Engelhard, B. et al. Specialized coding of sensory, motor and cognitive variables in VTA dopamine neurons. Nature 570, 509–513 (2019).

8. Nieh, E. H. et al. Geometry of abstract learned knowledge in the hippocampus. Nature 595, 80–84 (2021).

9. Bolkan, S. S. et al. Opponent control of behavior by dorsomedial striatal pathways depends on task demands and internal state. Nat. Neurosci. 25, 345–357 (2022).

10. Koay, S. A., Charles, A. S., Thiberge, S. Y., Brody, C. D. & Tank, D. W. Sequential and efficient neural-population coding of complex task information. Neuron 110, 328–349.e11 (2022).

11. Reiner, A. et al. Differential loss of striatal projection neurons in Huntington disease. Proc. Natl. Acad. Sci. 85, 5733–5737 (1988).

12. Krack, P. et al. Five-Year Follow-up of Bilateral Stimulation of the Subthalamic Nucleus in Advanced Parkinson’s Disease. N. Engl. J. Med. 349, 1925–1934 (2003).

13. Weaver, F. M. et al. Bilateral Deep Brain Stimulation vs Best Medical Therapy for Patients With Advanced Parkinson Disease: A Randomized Controlled Trial. JAMA 301, 63–73 (2009).

14. Eisinger, R. S., Cernera, S., Gittis, A., Gunduz, A. & Okun, M. S. A review of basal ganglia circuits and physiology: Application to deep brain stimulation. Parkinsonism Relat. Disord. 59, 9–20 (2019).

15. Mastro, K. J. et al. Cell-specific pallidal intervention induces long-lasting motor recovery in dopamine-depleted mice. Nat. Neurosci. 20, 815–823 (2017).

16. Spix, T. A. et al. Population-specific neuromodulation prolongs therapeutic benefits of deep brain stimulation. Science 374, 201–206 (2021).

17. Kravitz, A. V. et al. Regulation of parkinsonian motor behaviours by optogenetic control of basal ganglia circuitry. Nature 466, 622–626 (2010).

18. Roseberry, T. K. et al. Cell-Type-Specific Control of Brainstem Locomotor Circuits by Basal Ganglia. Cell 164, 526–537 (2016).

19. Bartholomew, R. A. et al. Striatonigral control of movement velocity in mice. Eur. J. Neurosci. 43, 1097–1110 (2016).

20. Yttri, E. A. & Dudman, J. T. Opponent and bidirectional control of movement velocity in the basal ganglia. Nature 533, 402–406 (2016).

21. Parker, J. G. et al. Diametric neural ensemble dynamics in parkinsonian and dyskinetic states. Nature 557, 177–182 (2018).

22. Bakhurin, K. I. et al. Opponent regulation of action performance and timing by striatonigral and striatopallidal pathways. eLife 9, e54831 (2020).

23. Lee, J., Wang, W. & Sabatini, B. L. Anatomically segregated basal ganglia pathways allow parallel behavioral modulation. Nat. Neurosci. 23, 1388–1398 (2020).

24. Chen, Z. et al. Direct and indirect pathway neurons in ventrolateral striatum differentially regulate licking movement and nigral responses. Cell Rep. 37, (2021).

25. Bonnavion, P. et al. Striatal projection neurons coexpressing dopamine D1 and D2 receptors modulate the motor function of D1- and D2-SPNs. Nat. Neurosci. 27, 1783–1793 (2024).

26. Cregg, J. M., Sidhu, S. K., Leiras, R. & Kiehn, O. Basal ganglia–spinal cord pathway that commands locomotor gait asymmetries in mice. Nat. Neurosci. 27, 716–727 (2024).

27. Balleine, B. W., Delgado, M. R. & Hikosaka, O. The Role of the Dorsal Striatum in Reward and Decision-Making. J. Neurosci. 27, 8161–8165 (2007).

28. Castañé, A., Theobald, D. E. H. & Robbins, T. W. Selective lesions of the dorsomedial striatum impair serial spatial reversal learning in rats. Behav. Brain Res. 210, 74–83 (2010).

29. Ding, L. & Gold, J. I. Separate, Causal Roles of the Caudate in Saccadic Choice and Execution in a Perceptual Decision Task. Neuron 75, 865–874 (2012).

30. Ding, L. & Gold, J. I. The basal ganglia’s contributions to perceptual decision-making. Neuron 79, 640–649 (2013).

31. Shadlen, M. N. & Shohamy, D. Decision Making and Sequential Sampling from Memory. Neuron 90, 927–939 (2016).

32. Yartsev, M. M., Hanks, T. D., Yoon, A. M. & Brody, C. D. Causal contribution and dynamical encoding in the striatum during evidence accumulation. eLife 7, e34929 (2018).

33. Malvaez, M. et al. Striatal cell-type specific stability and reorganization underlying agency and habit. 2025.01.26.634924 Preprint at 10.1101/2025.01.26.634924 (2025).

34. Collins, A. G. E. & Frank, M. J. Cognitive control over learning: Creating, clustering, and generalizing task-set structure. Psychol. Rev. 120, 190–229 (2013).

35. Hintiryan, H. et al. The mouse cortico-striatal projectome. Nat. Neurosci. 19, 1100–1114 (2016).

36. Hunnicutt, B. J. et al. A comprehensive excitatory input map of the striatum reveals novel functional organization. eLife 5, e19103 (2016).

37. Foster, N. N. et al. The mouse cortico–basal ganglia–thalamic network. Nature 598, 188–194 (2021).

38. Cuevas, N. et al. Context dependent contributions of the direct and indirect pathways in the associative and sensorimotor striatum. eLife 13, (2024).

39. Gradinaru, V. et al. Molecular and Cellular Approaches for Diversifying and Extending Optogenetics. Cell 141, 154–165 (2010).

40. Liu, L. D. et al. Accurate Localization of Linear Probe Electrode Arrays across Multiple Brains. eNeuro 8, (2021).

41. Laboratory, I. B. et al. Reproducibility of in vivo electrophysiological measurements in mice. eLife 13, (2024).

42. van Beest, E. H. et al. Tracking neurons across days with high-density probes. Nat. Methods 1–10 (2024) doi:10.1038/s41592-024-02440-1.

43. Pinto, L. et al. Task-Dependent Changes in the Large-Scale Dynamics and Necessity of Cortical Regions. Neuron 104, 810–824.e9 (2019).

44. Ashwood, Z. C. et al. Mice alternate between discrete strategies during perceptual decision-making. Nat. Neurosci. 25, 201–212 (2022).

45. Stujenske, J. M., Spellman, T. & Gordon, J. A. Modeling the Spatiotemporal Dynamics of Light and Heat Propagation for In Vivo Optogenetics. Cell Rep. 12, 525–534 (2015).

46. Owen, S. F., Liu, M. H. & Kreitzer, A. C. Thermal constraints on in vivo optogenetic manipulations. Nat. Neurosci. 22, 1061–1065 (2019).

47. Brown, L. S. et al. Neural circuit models for evidence accumulation through choice-selective sequences. bioRxiv 2023.09.01.555612 (2023) doi:10.1101/2023.09.01.555612.

48. Taverna, S., Ilijic, E. & Surmeier, D. J. Recurrent Collateral Connections of Striatal Medium Spiny Neurons Are Disrupted in Models of Parkinson’s Disease. J. Neurosci. 28, 5504–5512 (2008).

49. Burke, D. A., Rotstein, H. G. & Alvarez, V. A. Striatal Local Circuitry: A New Framework for Lateral Inhibition. Neuron 96, 267–284 (2017).

50. Assous, M. & Tepper, J. M. Excitatory extrinsic afferents to striatal interneurons and interactions with striatal microcircuitry. Eur. J. Neurosci. 49, 593–603 (2019).

51. Hjorth, J. J. J. et al. The microcircuits of striatum in silico. Proc. Natl. Acad. Sci. 117, 9554–9565 (2020).

52. Shi, W. et al. Whole-brain mapping of efferent projections of the anterior cingulate cortex in adult male mice. Mol. Pain 18, 17448069221094529 (2022).

53. Cox, J. et al. A neural substrate of sex-dependent modulation of motivation. Nat. Neurosci. 26, 274–284 (2023).

54. Gupta, D. et al. A multi-region recurrent circuit for evidence accumulation in rats. bioRxiv 2024.07.08.602544 (2024) doi:10.1101/2024.07.08.602544.

55. Bondy, A. G. et al. Coordinated cross-brain activity during accumulation of sensory evidence and decision commitment. 2024.08.21.609044 Preprint at 10.1101/2024.08.21.609044 (2024).

56. Wang, Y. et al. A cortico-basal ganglia-thalamo-cortical channel underlying short-term memory. Neuron 109, 3486–3499.e7 (2021).

57. Yang, W., Tipparaju, S. L., Chen, G. & Li, N. Thalamus-driven functional populations in frontal cortex support decision-making. Nat. Neurosci. 25, 1339–1352 (2022).

58. Ramot, A. et al. Motor learning refines thalamic influence on motor cortex. Nature 1–10 (2025) doi:10.1038/s41586-025-08962-8.

59. Phillips, J. M. et al. Primate thalamic nuclei select abstract rules and shape prefrontal dynamics. Neuron 113, 2014–2027.e12 (2025).

60. Rikhye, R. V., Gilra, A. & Halassa, M. M. Thalamic regulation of switching between cortical representations enables cognitive flexibility. Nat. Neurosci. 21, 1753–1763 (2018).

61. Khibnik, L. A., Tritsch, N. X. & Sabatini, B. L. A Direct Projection from Mouse Primary Visual Cortex to Dorsomedial Striatum. PLOS ONE 9, e104501 (2014).

62. Peters, A. J., Fabre, J. M. J., Steinmetz, N. A., Harris, K. D. & Carandini, M. Striatal activity topographically reflects cortical activity. Nature 591, 420–425 (2021).

63. Pan-Vazquez, A. et al. Pre-existing visual responses in a projection-defined dopamine population explain individual learning trajectories. Curr. Biol. 34, 5349–5358.e6 (2024).

64. Vu, M.-A. T. et al. Targeted micro-fiber arrays for measuring and manipulating localized multi-scale neural dynamics over large, deep brain volumes during behavior. Neuron 112, 909–923.e9 (2024).

65. Yin, H. H., Knowlton, B. J. & Balleine, B. W. Lesions of dorsolateral striatum preserve outcome expectancy but disrupt habit formation in instrumental learning. Eur. J. Neurosci. 19, 181–189 (2004).

66. Yin, H. H., Ostlund, S. B., Knowlton, B. J. & Balleine, B. W. The role of the dorsomedial striatum in instrumental conditioning. Eur. J. Neurosci. 22, 513–523 (2005).

67. Shiflett, M. W., Brown, R. A. & Balleine, B. W. Acquisition and Performance of Goal-Directed Instrumental Actions Depends on ERK Signaling in Distinct Regions of Dorsal Striatum in Rats. J. Neurosci. 30, 2951–2959 (2010).

68. Tai, L.-H., Lee, A. M., Benavidez, N., Bonci, A. & Wilbrecht, L. Transient stimulation of distinct subpopulations of striatal neurons mimics changes in action value. Nat. Neurosci. 15, 1281–1289 (2012).

69. Kravitz, A. V., Tye, L. D. & Kreitzer, A. C. Distinct roles for direct and indirect pathway striatal neurons in reinforcement. Nat. Neurosci. 15, 816–818 (2012).

70. Hikosaka, O., Kim, H. F., Yasuda, M. & Yamamoto, S. Basal ganglia circuits for reward value-guided behavior. Annu. Rev. Neurosci. 37, 289–306 (2014).

71. Nonomura, S. et al. Monitoring and Updating of Action Selection for Goal-Directed Behavior through the Striatal Direct and Indirect Pathways. Neuron 99, 1302–1314.e5 (2018).

72. Kwak, S. & Jung, M. W. Distinct roles of striatal direct and indirect pathways in value-based decision making. eLife 8, e46050 (2019).

73. Peak, J., Chieng, B., Hart, G. & Balleine, B. W. Striatal direct and indirect pathway neurons differentially control the encoding and updating of goal-directed learning. eLife 9, e58544 (2020).

74. Jang, H. J., Ward, R. M., Golden, C. E. M. & Constantinople, C. M. Acetylcholine demixes heterogeneous dopamine signals for learning and moving. bioRxiv 2024.05.03.592444 (2024) doi:10.1101/2024.05.03.592444.

75. Reinhold, K. et al. Striatum supports fast learning but not memory recall. Nature 1–10 (2025) doi:10.1038/s41586-025-08969-1.

76. Thorn, C. A., Atallah, H., Howe, M. & Graybiel, A. M. Differential Dynamics of Activity Changes in Dorsolateral and Dorsomedial Striatal Loops During Learning. Neuron 66, 781–795 (2010).

77. Gremel, C. M. & Costa, R. M. Orbitofrontal and striatal circuits dynamically encode the shift between goal-directed and habitual actions. Nat. Commun. 4, 2264 (2013).

78. Graybiel, A. M. Habits, Rituals, and the Evaluative Brain. Annu. Rev. Neurosci. 31, 359–387 (2008).

79. Balleine, B. W. & O’Doherty, J. P. Human and Rodent Homologies in Action Control: Corticostriatal Determinants of Goal-Directed and Habitual Action. Neuropsychopharmacology 35, 48–69 (2010).

80. Dolan, R. J. & Dayan, P. Goals and Habits in the Brain. Neuron 80, 312–325 (2013).

81. Mizes, K. G. C., Lindsey, J., Escola, G. S. & Ölveczky, B. P. Dissociating the contributions of sensorimotor striatum to automatic and visually guided motor sequences. Nat. Neurosci. 26, 1791–1804 (2023).

82. Tsutsui-Kimura, I. et al. Dopamine in the tail of the striatum facilitates avoidance in threat–reward conflicts. Nat. Neurosci. 1–16 (2025) doi:10.1038/s41593-025-01902-9.

83. Lowet, A. S. et al. An opponent striatal circuit for distributional reinforcement learning. Nature 639, 717–726 (2025).

84. Markowitz, J. E. et al. The Striatum Organizes 3D Behavior via Moment-to-Moment Action Selection. Cell 174, 44–58.e17 (2018).

85. Dhawale, A. K., Wolff, S. B. E., Ko, R. & Ölveczky, B. P. The basal ganglia control the detailed kinematics of learned motor skills. Nat. Neurosci. 24, 1256–1269 (2021).

86. Tang, Y. et al. Opposing regulation of short-term memory by basal ganglia direct and indirect pathways that are coactive during behavior. 2021.12.15.472735 Preprint at 10.1101/2021.12.15.472735 (2021).

87. Luo, T. Z. et al. An approach for long-term, multi-probe Neuropixels recordings in unrestrained rats. eLife 9, e59716 (2020).

88. Steinmetz, N. A. et al. Neuropixels 2.0: A miniaturized high-density probe for stable, long-term brain recordings. Science (2021) doi:10.1126/science.abf4588.

89. Aronov, D. & Tank, D. W. Engagement of neural circuits underlying 2D spatial navigation in a rodent virtual reality system. Neuron 84, 442–456 (2014).

90. Hanks, T. D. et al. Distinct relationships of parietal and prefrontal cortices to evidence accumulation. Nature 520, 220–223 (2015).

91. Pisanello, F. et al. Dynamic illumination of spatially restricted or large brain volumes via a single tapered optical fiber. Nat. Neurosci. 20, 1180–1188 (2017).

92. Zimmerman, C. A. et al. A neural mechanism for learning from delayed postingestive feedback. Nature 1–10 (2025) doi:10.1038/s41586-025-08828-z.

93. Pachitariu, M., Sridhar, S. & Stringer, C. Solving the spike sorting problem with Kilosort. bioRxiv 2023.01.07.523036 (2023) doi:10.1101/2023.01.07.523036.

94. Findling, C. et al. Brain-wide representations of prior information in mouse decision-making. 2023.07.04.547684 Preprint at 10.1101/2023.07.04.547684 (2024).

95. Laboratory, I. B. et al. A Brain-Wide Map of Neural Activity during Complex Behaviour. 2023.07.04.547681 Preprint at 10.1101/2023.07.04.547681 (2024).

96. Inagaki, H. K. et al. A midbrain-thalamus-cortex circuit reorganizes cortical dynamics to initiate movement. Cell 185, 1065–1081.e23 (2022).

97. Scott, B. B. et al. Fronto-parietal Cortical Circuits Encode Accumulated Evidence with a Diversity of Timescales. Neuron 95, 385–398.e5 (2017).

98. Brunton, B. W., Botvinick, M. M. & Brody, C. D. Rats and humans can optimally accumulate evidence for decision-making. Science 340, 95–98 (2013).

99. Renier, N. et al. iDISCO: A Simple, Rapid Method to Immunolabel Large Tissue Samples for Volume Imaging. Cell 159, 896–910 (2014).

100. Renier, N. et al. Mapping of Brain Activity by Automated Volume Analysis of Immediate Early Genes. Cell 165, 1789–1802 (2016).

101. Pisano, T. J. et al. Homologous organization of cerebellar pathways to sensory, motor, and associative forebrain. Cell Rep. 36, (2021).

102. Dean, K. M., Roudot, P., Welf, E. S., Danuser, G. & Fiolka, R. Deconvolution-free Subcellular Imaging with Axially Swept Light Sheet Microscopy. Biophys. J. 108, 2807–2815 (2015).

103. Bria, A. & Iannello, G. TeraStitcher - A tool for fast automatic 3D-stitching of teravoxel-sized microscopy images. BMC Bioinformatics 13, 316 (2012).

104. Swaney, J. et al. Scalable image processing techniques for quantitative analysis of volumetric biological images from light-sheet microscopy. 576595 Preprint at 10.1101/576595 (2019).

105. Tyson, A. L. et al. Accurate determination of marker location within whole-brain microscopy images. Sci. Rep. 12, 867 (2022).

